# *In vitro* neurons learn and exhibit sentience when embodied in a simulated game-world

**DOI:** 10.1101/2021.12.02.471005

**Authors:** Brett J. Kagan, Andy C. Kitchen, Nhi T. Tran, Bradyn J. Parker, Anjali Bhat, Ben Rollo, Adeel Razi, Karl J. Friston

## Abstract

Integrating neurons into digital systems to leverage their innate intelligence may enable performance infeasible with silicon alone, along with providing insight into the cellular origin of intelligence. We developed *DishBrain*, a system which exhibits natural intelligence by harnessing the inherent adaptive computation of neurons in a structured environment. *In vitro* neural networks from human or rodent origins, are integrated with *in silico* computing via high-density multielectrode array. Through electrophysiological stimulation and recording, cultures were embedded in a simulated game-world, mimicking the arcade game ‘Pong’. Applying a previously untestable theory of active inference via the Free Energy Principle, we found that learning was apparent within five minutes of real-time gameplay, not observed in control conditions. Further experiments demonstrate the importance of closed-loop structured feedback in eliciting learning over time. Cultures display the ability to self-organise in a goal-directed manner in response to sparse sensory information about the consequences of their actions.

---

Harnessing the computational power of living neurons to create synthetic biological intelligence (SBI), previously confined to the realm of science fiction, is now tantalisingly within the reach of human innovation. The superiority of biological computation has been widely recognised with attempts to develop hardware supporting neuromorphic computing^1^. Yet, no system outside biological neurons are capable of supporting at least third-order complexity which is necessary to recreate the complexity of a biological neuronal network (BNN)^1,2^. This raises significant challenges to any attempts to generate *in silico* neuronal models to predict function of BNN systems^3^. Here we aim to establish functional *in vitro* networks of cortical cells from embryonic rodent and human induced pluripotent stem cells (hiPSCs) on high-density multielectrode arrays (HD-MEA) to demonstrate that these neural cultures can exhibit biological intelligence—as evidenced by learning in a simulated gameplay environment—in real time (Figure 1). Being able to successfully interact with SBIs would enable investigations into previously untestable areas. This would include, but not limited to, pseudo-cognitive responses as part of drug screening, bridging the divide between single cell and population coding approaches to understanding neurobiology, better understanding how BNNs compute to inform machine learning approaches, and potentially give rise to silico-biological computational platforms that surpass the performance of existing silicon-alone hardware. Indeed, some proponents suggest that generalised SBI may arrive before artificial general intelligence (AGI) due to the inherent efficiency and evolutionary advantage of biological systems^4^.

**Fig. 1.**
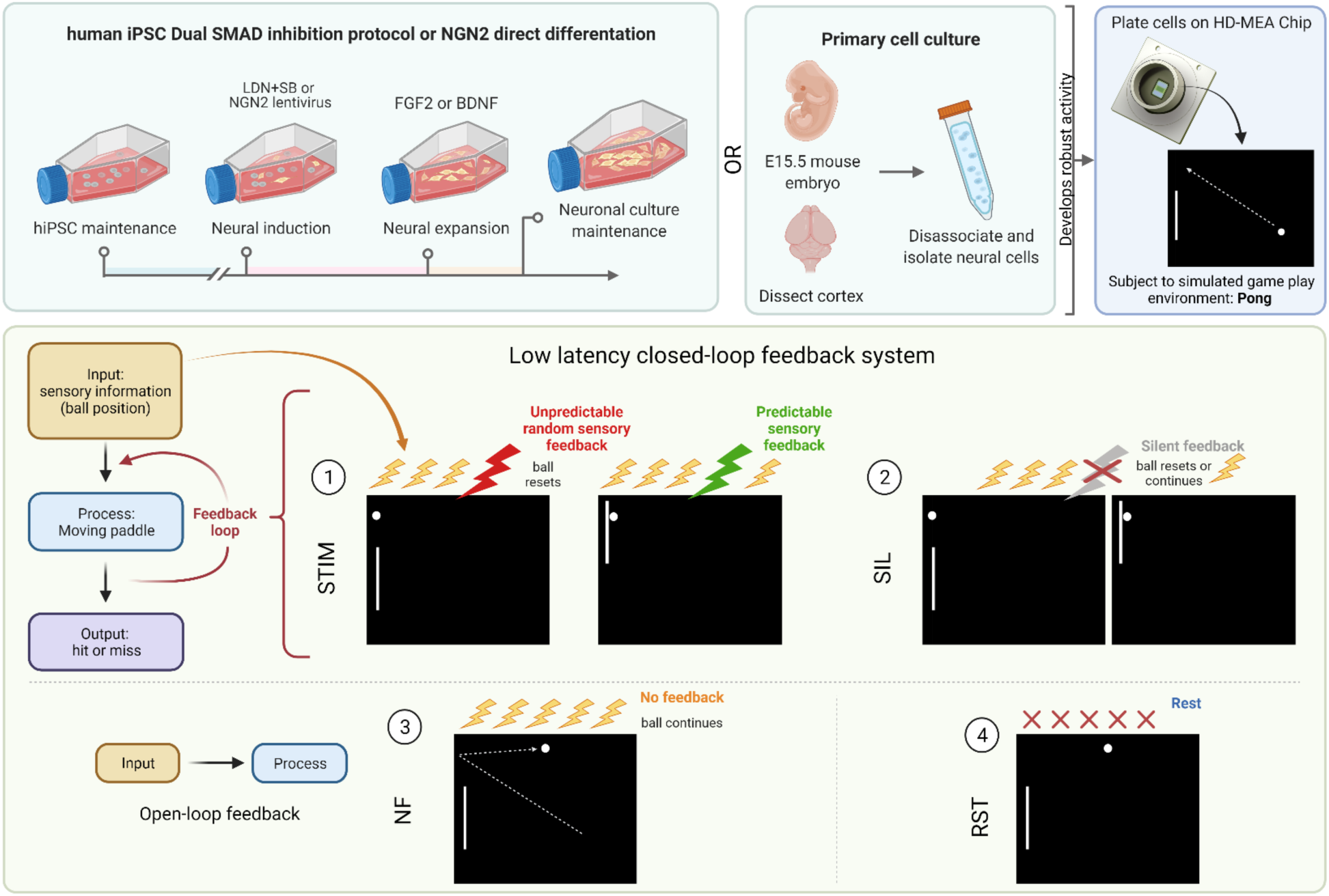
*DishBrain* system and experimental protocol schematic. Neuronal cultures derived from either human induced pluripotent stem cells (iPSC) via Dual SMAD inhibition, NGN2 lentivirus directed differentiation, or primary cortical cells from E15.5 mouse embryos, were plated onto HD-MEA chips and embedded in a stimulated game-world of ‘pong’ via the *DishBrain* system. Different *DishBrain* environments were utilised to demonstrate: (1&2) low latency closed-loop feedback system (stimulation (STIM) & silent (SIL) treatment); (3) No feedback (NF) system to demonstrate an open-loop feedback configuration; and (4) rest (RST) configuration to demonstrate a system in which sensory information (yellow bolt) is absent. An interactive visualiser with gameplay is available at https://bit.ly/3DSi4Eg

This system that we termed *DishBrain*, can leverage the inherent property of neurons to share a ‘language’ of electrical (synaptic) activity with each other to link silicon and BNN systems through electrical stimulation and recording. Given the compatibility of hardware and cells, wetware, there are two interrelated processes that are required for sentient behaviour in an intelligent system. Firstly, the system must learn how external states influence internal states—via perception—and how internal states influence external states—via action. Secondly, the system must infer from its sensory states when it should adopt a particular behaviour. In short, it must be able to predict how its actions will influence the environment. To address the first imperative, custom software drivers were developed to create low latency closed-loop feedback systems that simulated exchange with an environment for BNNs through electrical stimulation. Closed-loop systems afford an *in vitro* culture ‘embodiment’ by providing feedback on the causal effect of the behaviour from the cell culture. Embodiment requires a separation of internal *vs* external states, where feedback of the effect of action on a given environment is available. Previous works both *in vitro* and *in silico* have shown that electrophysiological closed-loop feedback systems engender significant network plasticity and potentially behavioural adaptation over and beyond what can be achieved with open-loop systems^5,6^. Further support for the link between embodiment and functional behaviour is found *in vivo* where disrupting a closed loop system by uncoupling visual feedback and motor outputs disrupts functional development of visual processing in the primary visual cortex in mice^7^. This strongly supports the vital link between feedback and the eventual development of functional behaviour in biological neural networks.

To address the second requirement, the system can be used to test key theories for how intelligent behaviour may arise. One proposition for how intelligent behaviour may arise in an intelligent system embodied in an environment is found in the theory of active inference via the Free Energy Principle (FEP)^8^. Previous work has established that neurons can perform blind-source separation via a state-dependent Hebbian plasticity that is consistent with the FEP^9,10^. The FEP suggests that any self-organising system separate from its environment seeks to minimizes its variational free energy^11–13^. This means that systems like the brain—at every spatiotemporal scale— may engage in active inference by using an internal generative model to predict sensory inputs that represent the external world^11–13^. The gap between the model predictions and observed sensations (‘surprise’ or ’prediction error’) may be minimised in two ways: by optimising probabilistic (Bayesian) beliefs about the environment to make predictions more like sensations, or by *acting* upon the environment to make sensations conform to its predictions. This implies a common objective function for action and perception that scores the fit between an internal model and the external environment.

Under this theory, BNNs hold ‘beliefs’ about the state of the world, where learning involves updating these beliefs to minimise their variational free energy or change the world, by action, to make it less surprising^13,14^. If true, this implies that it should be possible to shape BNN behaviour by simply presenting noisy, unpredictable feedback following ’incorrect’ behaviour. If BNNs are presented with unpredictable feedback, they should adopt actions that avoid the states that resulted in this input. By developing a system that allows for neural cultures to be embodied in a simulated game-world, we are not only able to test whether these cells are capable of engaging in goal-directed learning in a dynamic envrionment, we are able to investigate a fundemental basis of intelligence.

## RESULTS

### Growth of neuronal ‘wetware’ for computation

Neurons can be grown or harvested in numerous ways. Cortical cells from the dissected cortices of rodent embryos can be grown on MEA in nutrient rich medium and maintained for months^15,16^. These cultures will develop complicated morphology, with numerous dendritic and axonal connections, leading to functional BNNs^17,18^. We successfully replicated the development of these cultures from embryonic day 15.5 (E15.5) mouse embryos, with a representative culture shown in Figure 2A. We also differentiated human induced pluripotent stem cells (hiPSCs) into monolayers of active heterogeneous cortical neurons which have been shown to display mature functional properties^19–21^. Using a dual SMAD inhibition as previously described^21,22^ we developed long-term cortical neurons that formed dense connections with supporting glial cells (Figure 2B - 2C). Finally, we wished to expand our study using a different method of hiPSC differentiation – NGN2 direct reprogramming – used in our final part of this study. Previous work has shown that human fibroblasts can be directly converted into induced neuronal cells which express a cortical phenotype^23,24^. This high yield method was replicated in this work with cells displaying pan-neuronal markers (Figure S1A, S1B). These cells typically display a high proportion of excitatory glutamatergic cells, quantified using qPCR shown in Figure 1D. Integration of these cells on the HD-MEA was confirmed via scanning electron microscopy (SEM) where cells had been maintained for > 3 months. Routine SEM imaging revealed dense clustering of neurons, with a clear contrast between cell and the MEA surface (Figure 2E). Densely interconnected dendritic networks could be observed in neuronal cultures forming interlaced networks spanning the MEA area (Figure 2F). These neuronal cultures appeared to rarely follow the topography of the MEA and were more likely to form large clusters of connected cells with dense dendritic networks (Figure 2G, 2H). This is likely due to the large size of an individual electrode within the MEA, however, there are also chemotactic effects that can contribute to counteract the effect of substrate topography on neurite projections^25^.

**Fig. 2.**
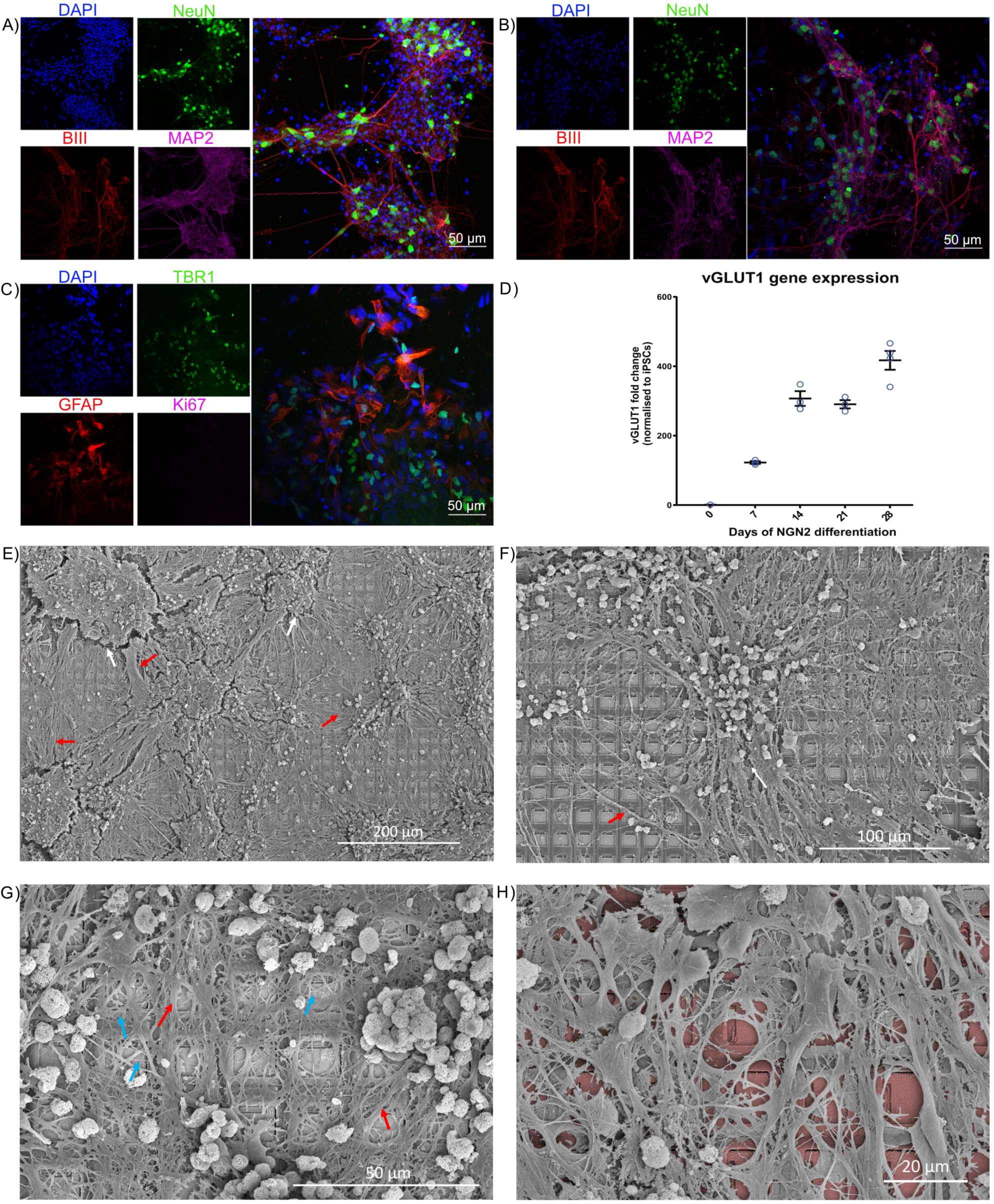
Cortical cells form dense interconnected networks. Scale bars as shown on panel. **A)** and **B)** show cortical cells harvested from embryonic rodents and differentiated from hIPSCs respectively. DAPI in blue stains all cells, NeuN in green shows neurons, BIII marks axons, while MAP2 marks dendrites. Further characterisation in **C)** with GFAP shows supporting astrocyte cells, critical for long-term functioning, along with a marker for cortex specific cells, TBR1. A risk with using IPSCs is that cells are not fully specified and may aggressively continue dividing, staining for Ki67, a marker of dividing cells, shows this is not a concern with these cultures. **D)** Gene expression studies over 28 days demonstrated increased expression of the glutamatergic neural marker, vesicular glutamate transporter 1 (vGLUT1). This data demonstrates that cells produced by NGN2 differentiation are comprised of synaptically active excitatory neural cells. **E) – G)** Neurons maintained on MEA for > 3 months. White arrows show regions of shrinkage within the cultures, red arrows show bundles of axons, blue arrays show single neurite extensions. Note complex and extensive connections between cells, the dense coverage over HD-MEA, and overlapping connections extended from neuronal soma present in all cultures, showing that cells overlap multiple electrodes. **H)** has false colouring to highlight the HD-MEA electrodes beneath the cells.

### Neural cells show well-characterised spontaneous action potentials which develop over time

We mapped the *in vitro* development of electrophysiological activity in neural systems at high spatial and temporal resolution. Robust activity in primary cortical cells from E15.5 rodents was found at days *in vitro* (DIV)14 (Figure 3A, 3E) where bursts of synchronised activity was regularly observed as previously demonstrated^17,18^. In contrast, while similar to previous reports, synchronised bursting activity was not observed in cortical cells from an hiPSC background differentiated using the dual SMAD inhibition protocol (DSI) until DIV 73 (Figure 3A, 3F)^19^. hiPSCs differentiated using NGN2 direct reprogramming showed activity much earlier, typically between days 14 and 24 (Figure 3A, 3G). Daily activity scans of electrophysiological activity were also conducted. While max firing rate typically increased and remained relatively stable over time for all cell types during the testing period (Figure 3B), changes were observed in both the mean firing rate (Figure 3C) and variance in firing rate (Figure 3D) over the days of testing. In particular, hiPSCs differentiated using the NGN2 direct reprogramming method showed a considerable increase in mean firing rate and the variance in firing over days.

**Fig. 3.**
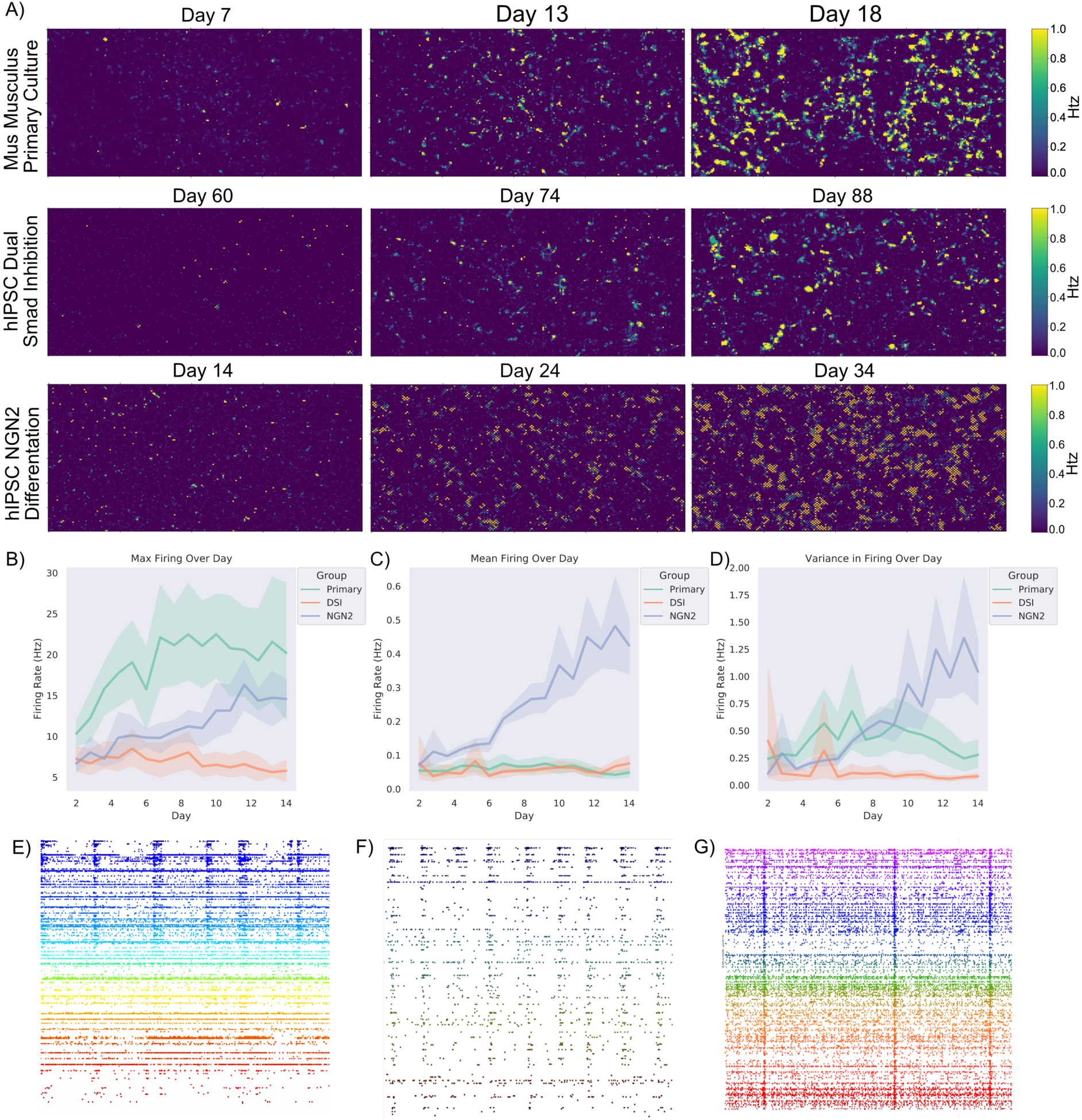
Cortical cells display spontaneous electrophysiological activity. Shaded error = 95% confidence intervals. **A)** Scale bar on the right indicates the frequency of firing in Hz. Displays the rate of firing over a representative culture grown from E15.5 primary rodent cortical cells, hIPSC cells differentiated to cortical neurons via dual SMAD inhibition (DSI), and hIPSC cells differentiated to cortical cells via NGN2 direct differentiation. Note that while all cultures show substantial firing over the majority of the assay area, they do so at different timepoints. Training was started when cells displayed consistent firing with a mean above 0.7Hz and continued over approximately 14 days, as seen in **B)** the max firing remained consistently different between cortical cells from a primary source and cortical cells differentiated from hIPSCs. Of interest though as seen in **C)** is that the mean activity between hIPSCs differentiated using DSI and primary cortical cultures was generally similar, while hIPSCs differentiated using the NGN2 method continued to increase. This is reflected in **D)** where the former two cell types displayed minimal changes in the variance in firing within a culture, while the latter increased variance over time. **E)**, **F)** & **G)** Showcases raster-plots over 50 seconds, where each dot is a neuron firing an action potential. Note the differences between mid-stage cortical cells from a DIV14 primary rodent culture (**E**) compared to more mature DIV73 human cortical cells (**F**) differentiated from iPSCs using the dual SMAD inhibition and NGN2 direct differentiated neurons (**G**) approach described in text, in terms of synchronised activity and stable firing patterns. While all display synchronised activity, there is a difference in the overall levels of activity represented in **B** - **D**.

### Building a modular, real-time platform to harness neuronal computation

We developed the *DishBrain* system to leverage neuronal computation and interact with neurons in an embodied environment (**Supp. Text 1**; Figure 4A). The *DishBrain* environment is a low latency, real-time system that interacts with the vendor MaxOne software, allowing it to be used in ways that extend its original functions (Figure 4B). This system can record electrical activity in a neuronal culture and provide external (non-invasive) electrical stimulation in a comparable manner to the generation of action potentials by internal electrical stimulation^26^. Using the coding schemes described in methods, external electrical stimulations convey a range of information: predictable, random, or sensory (Figure 5A). This setup enables one to not only ‘read’ information from a neural culture, but to ‘write’ sensory data into one. The initial proof of principle using *DishBrain* was to simulate the classic arcade game ‘pong’ by delivering inputs to a predefined sensory area. Similarly, the electrophysiological activity of pre-defined motor regions was gathered—in real time—to move a ‘paddle’. Preliminary investigations compared different motor region configurations using an EXP3 algorithm (**Supp. Text 2**; **Figure S3**). This aimed to identify whether the neural cultures had activity that was more successful under specific configurations by choosing setups that resulted in a higher hit rate. Experimental cultures showed significantly different preferences for configurations compared to media-only controls (Figure 5B). While media-only controls showed a preference for configurations that maximised bias—where sensory stimulation alone could direct the gameplay towards higher performance (blinded fully later)—experimental cultures showed a preference for configuration that enabled lateral inhibition (Figure 5C).

**Fig. 4.**
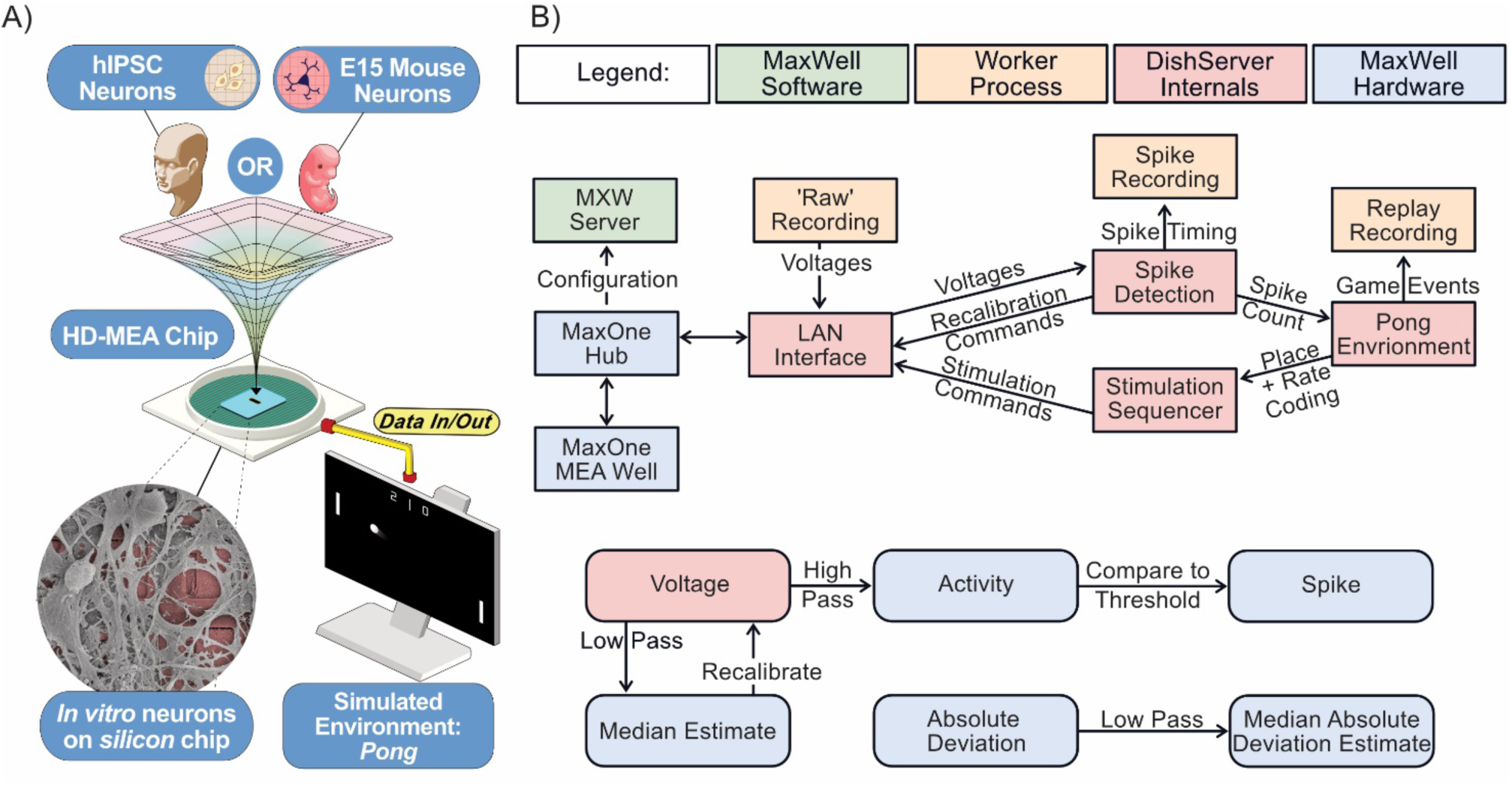
Schematics of software used for *DishBrain*. **A)** Diagrammatic overview of DishBrain setup. **B)** Software components and data flow in the DishBrain closed loop system. Voltage samples flow from the MEA to the ‘pong’ environment, and sensory information flows back to the MEA, forming a closed loop. The blue rectangles mark proprietary pieces of hardware from MaxWell while the green MXWServer is used to configure the MEA and Hub. Red rectangles mark components of the ’DishServer’ program, a high-performance program consisting of four components designed to run asynchronously. Running a virtual environment in a closed loop imposes strict performance requirements, and digital signal processing is the main bottleneck of this system. The ’LAN Interface’ component stores network state, for talking to the Hub, and produces arrays of voltage values for processing. Voltage values are passed to the ’Spike Detection’ component, which stores feedback values and spike counts, and passes recalibration commands back to the LAN Interface. When the pong environment is ready to run, it updates the state of the paddle based on the spike counts, updates the state of the ball based on its velocity and collision conditions, and reconfigures the stimulation sequencer based on the relative position of the ball and current state of the game. The stimulation sequencer stores and updates indices and countdowns relating to the stimulations it must produce and converts these into commands each time the corresponding countdown reaches zero, which are finally passed back to the LAN Interface, to send to the MEA system, closing the loop. Numeric operations in the real-time spike detection component of the *DishBrain* closed loop system are shown below, including multiple IIR filters.

**Fig. 5.**
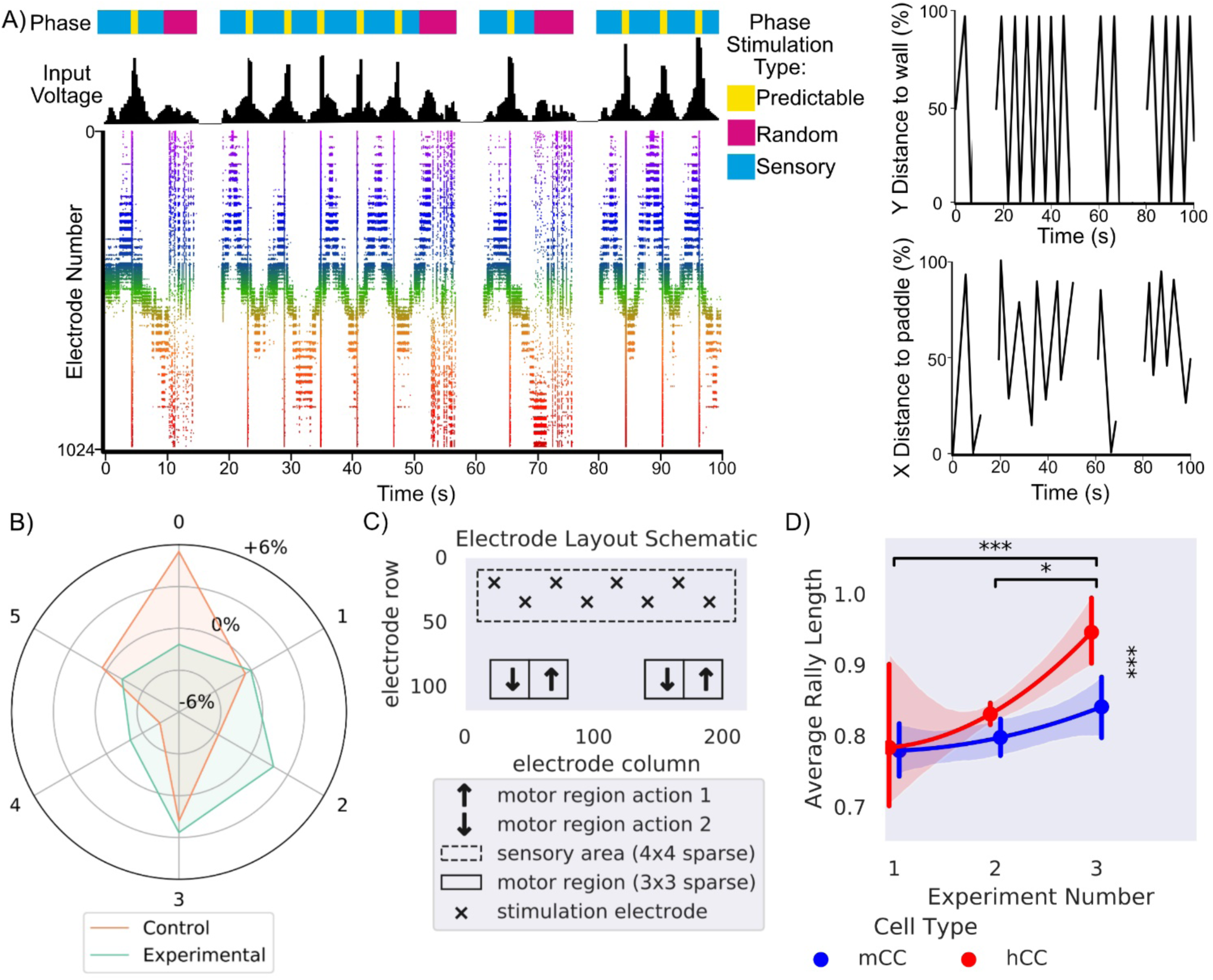
Schematics, EXP3 configuration selection and testing with increasing informational density. **A)** Presents a schematic showing the different phases of stimulation which provides information about the environment to the culture, in line with this is the corresponding input voltage and how that voltage appears on the raster plot over 100 seconds. The appearance of random stimulation after a ball missing vs system wide predictable stimulation upon a successful hit is apparent across all three representations. This corresponds to the images on the right which show the position of the ball on both x and y axis relative to the paddle and backwall in % of total distance is shown on the same timescale. **B)** Shows the distribution differences relative to chance in percentage that a motor configuration was chosen by EXP3 algorithm (*χ2* = 35690.93, *p*<0.0001) for control and experimental cultures. Motor configuration 0 was selected most often for media control while motor configuration 3 was selected most often for experimental. **C)** Final electrode layout schematic for *DishBrain* pong-world gameplay. **D)** * = *p* < 0.05, *** = *p* <0.001, Error bars = 95% CI, shows average rally length over three distinct experiment rounds during design of *DishBrain* pong-world where each subsequent experiment provided higher density information on ball position than the previous.

### Increasing the density of sensory information input leads to increased performance

The *DishBrain* protocol was refined over a series of pilot studies, which can be grouped into three broad experiments, each increasing in density of sensory information. The first experiment operated with a 4Hz stimulation that was purely rate coded. Experiment two included the EXP3 based configuration. Experiment 3 removed the EXP3 based configuration, locked to the layout in Figure 5C, and change to the combined rate (4 – 40Hz) and placed coding method of data input. Notably the biggest increase was between second and third experiments with the introduction of rate coding of the ball’s position to supplement the purely rate coded approach used previously. The gameplay for the final fifteen minutes for each culture type was compared (Figure 5D; Table S1). Cultures displayed a significant increase in performance between the second and final session and the first and final session. Between cultures, human cortical cells (hCCs) had significantly longer average rally lengths than cultures with mice cortical cells (mCCs)(Table S2). This is interesting because it suggests that—at a neuronal level—cortical cells from a human origin, in this case from hiPSCs, can out-perform cells from an embryonic mouse, even when overall cell numbers are comparable. Overall, the magnitude of this change supports that increasing sensory information successively improved performance, even when cell culture features were kept constant.

### Biological neural networks learn over time when embodied in a gameplay environment

To test the theory of active inference via the FEP (Figure 6A), using the parameters described in **Methods**, cortical cells, mCC and hCCs, were compared to media-only controls (CTL), rest sessions—where active cultures controlled the paddle but received no sensory information (RST), and to in-silico controls that mimicked all aspect of the gameplay but paddle was driven by random noise (IS), over 399 test-sessions (80-CTL, 42-RST, 38-IS, 101-mCCs, 138-hCCs). The average rally length (total number of successful ball intercepts by the paddle) showed a significant interaction (Figure 6B; Table S1), with differences occurring based on a combination of group and time (first five and last fifteen minutes). Only the mCC and hCC cultures showed evidence of learning over time, with significantly increased rally lengths at the second timepoint compared to the first. Further, it was found that during the first five minutes of gameplay key significant differences were observed (Table S1). The hCC group performed significantly worse than mCC, CTL and IS groups (Table S2). This suggests that hCCs seem to perform worse than controls when first embodied in an environment, suggesting an initially maladaptive control of the paddle. Notably, at the latter timepoint this trend was reversed, the hCC group significantly outperformed all control groups along with a slight but significant difference over the mCC group (Table S1). Likewise, the mCC group significantly outperformed all control groups (Table S2). This data replicates our earlier finding on the differences between mouse and human cells, along with unambiguously demonstrating a significant learning effect in both experimental groups that was absent in the control groups (**Movie S1**).

**Fig. 6.**
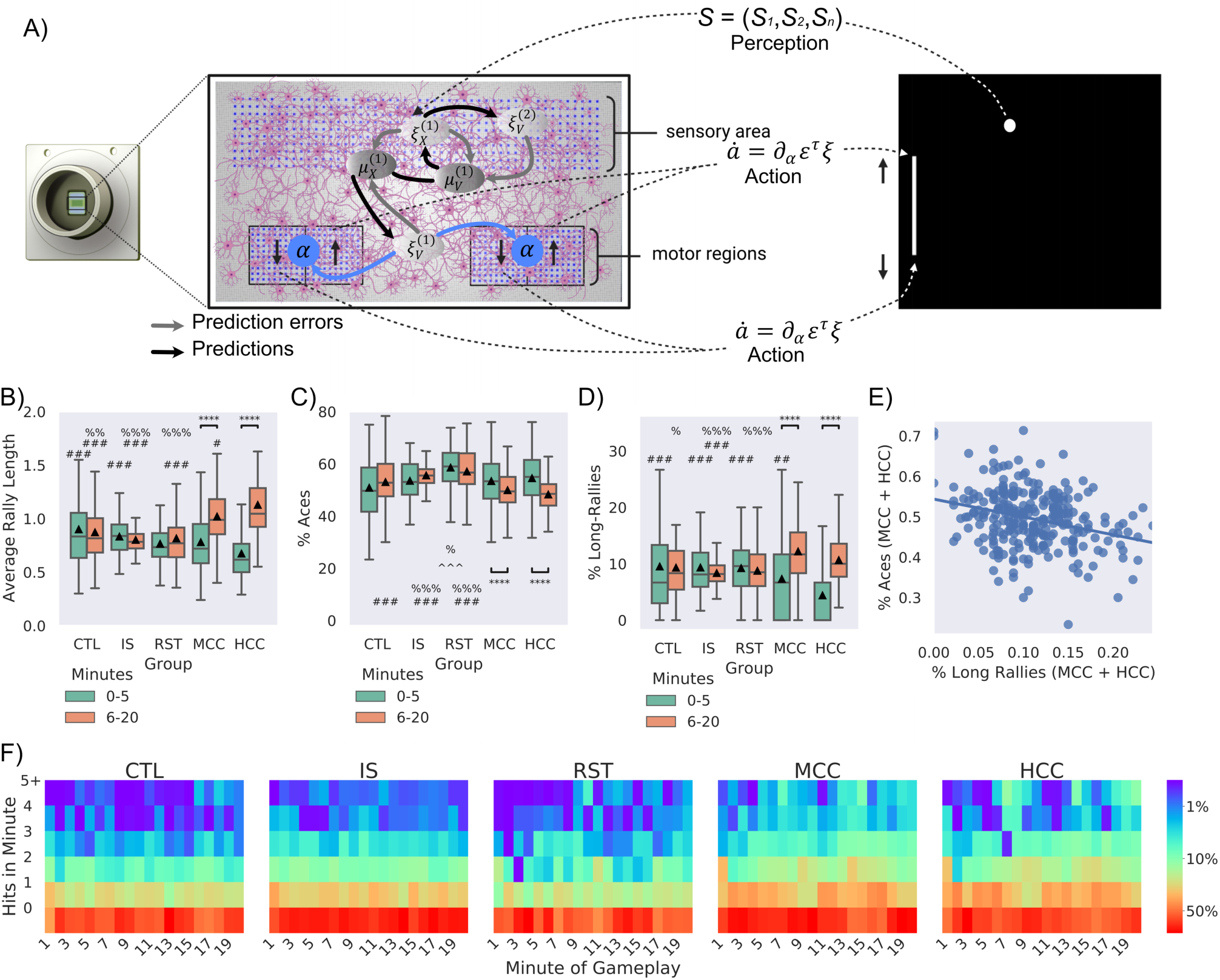
Embodied Cortical Neurons Show Significantly Improved Performance in Pong When Embodied in a Virtual Game-World. Significance bars show within group differences denoted with *. Symbols show between group differences at the given timepoint: # = vs HCC; % = *vs* MCC; ^^ = vs CTL; @ = vs IS. The number of symbols denotes the p-value cut off, where 1 = p < 0.05, 2 = p <0.01, 3 = p < 0.001 and 4 = p <0.0001. Box plots show interquartile range, with bars demonstrating 1.5X interquartile range, the line marks the median and ▴ marks the mean. **A)** Schematic of how neurons may engage in the game-world under active inference. This illustration adopts a predictive coding (a.k.a. Kalman filter) formulation of variational free energy minimisation, in which neuronal dynamics are read as gradient flows—and free energy gradients are read as prediction errors. On this view, prediction errors can be regarded as driving neuronal activity—that implicitly parameterises a generative or forward model—and motor responses, via minimising (synthetic) proprioceptive prediction errors. **B-D)** compares experimental groups according to two time points: timepoint 1: first 5 mins of gameplay (0-5 mins), timepoint 2: last 15 mins of gameplay (6-20 mins). **B)** Average performance between groups over time, where only experimental (MCC: *t* = 6.15, *p* = 5.27^-08^ & HCC: *t* = 10.44, *p* = 3.91^-19^) showed significant improvement and higher performance against all control groups at the second timepoint. **C)** Average number of aces between groups and over time, only MCC (*t* = 2.67, *p* = 0.008) and HCC (*t* = 5.95, *p* = 2.13^-08^) differed significantly over time. The RST group had significantly more aces compared to the CTL, IS, MCC, and HCC groups at timepoint one and compared to the CTL, MCC and HCC at timepoint 2. Only MCC and HCC showed significant decreases in the number of aces over time, indicating learning. At the latter timepoint they also showed fewer aces compared to the IS group, but only the HCC group was significantly less than CTL. **D)** Average number of long-rallies (>3) performed in a session. At timepoint 1, the HCC group had significantly fewer long-rallies compared to all control groups (CTL, IS, and RST). However, both the MCC (*t* = 5.55, *p* = 2.36^-07^) and HCC (*t* = 10.38, *p* = 5.27^-19^) groups showed significantly more long-rallies over time. As such, by timepoint 2, the HCC group displayed significantly more long-rallies compared to the IS group. The HCC group also displayed significantly more long-rallies compared to all CTL, IS, and RST control groups. **E)** Significant negative correlation (*r* = -0.34, *p* < 0.001) between % aces and % long rallies for experimental cultures in the last 15 minutes. **F)** Distribution of frequency of mean summed hits per minute amongst groups show obvious differences.

### Nuances between how learning occurs exhibits differences between cell types

To determine how the above learning arose, key gameplay characteristics were examined further. The number of times the paddle failed to intercept the ball on the initial serve (aces; Figure 6C) and the number of long rallies (> 3 consecutive hits; Figure 6D) were calculated for this data. As with average rally length, significant interactions between groups and time were found both for aces and long rallies (Table S1). Only the mCC and hCC groups showed significantly fewer aces in the latter timepoint compared to the first (Table S2). Likewise, only the mCC and hCC groups showed significantly more long rallies in the latter timepoint compared to the first (Table S2). This shows that both experimental cultures improved performance by not only reducing how often they missed the initial serve, but by achieving more consecutive hits also. Similarly, a significant difference between groups was found both for aces and long rallies (Table S1). At the first timepoint for aces, it was found that the RST condition had significantly more aces than the CTL and mCC groups (Table S2). It is difficult to determine exactly why allowing the paddle to be controlled by unstimulated cells would result in more aces initially than other groups. Perhaps there is a degree of sporadic behaviour that the cells engage with when initially introduced to the rest period from gameplay that results in this behaviour, possibly similar to what was observed in average rally length by hCCs above. When the number of long-rallies at this time point was investigated, it was found that only HCC had significantly fewer long-rallies compared to all groups (Table S2). This is consistent with the finding that hCC do show worse performance in the first time point overall and explains why this may be observed.

Significant differences between groups at the latter timepoint was also found both for aces and long rallies (Table S1). Most notably the HCC group showed significantly fewer aces compared to CTL, RST, and IS groups (Table S1). The mCC group also showed significantly fewer aces compared to RST and IS groups, however not the CTL group (Table S2). In contrast, for long-rallies the mCC group showed significantly more than the CTL, RST and IS groups (Table S2). Yet the hCC group only showed significantly more long-rallies compared to the IS group, but not RST or CTL (Table S2). Moreover, here a significant negative correlation was found, suggesting that the performance was not arising out of a maladaptive behaviour such as fixing the paddle to a single corner (Figure 6E). Wholistically, Figure 6F emphasises that although both mCC and hCC showed fewer aces and more long-rallies in the latter timepoints compared to the first, the cell types did display nuances in their behaviour, highlighting differences between cell types. The data also further suggests that an unstimulated culture still controlling the paddle will have significantly poorer performance than controls where the paddle is moved based on noise. This does suggest a systematic control that is difficult to interpret from this data but does indicate the potential of an enduring embodiment once stimulation ceases.

### Biological neural networks require feedback for learning

To investigate the importance of feedback type for learning, cultures were tested under three conditions, for three days, with three sessions per day resulting in 483 sessions. Condition 1 (Stimulus) mimicked that used above, where predictable and unpredictable stimuli were administered when the cultures behaved desirably or not, respectively. Condition 2 (Silent) involved the stimulus feedback being replaced with a matching time-period in which all stimulation was withheld. Condition 3 (No-feedback) removed the restart after a miss. When the paddle did not successfully intercept the ball, the ball would bounce and continue without interruption: the stimulus reporting ball position was still provided. The difference between these conditions is emphasised in Figure 7A. Rest period activity was also gathered used to normalise performance on a per session basis to account for differences in unstimulated activity.

**Fig 7.**
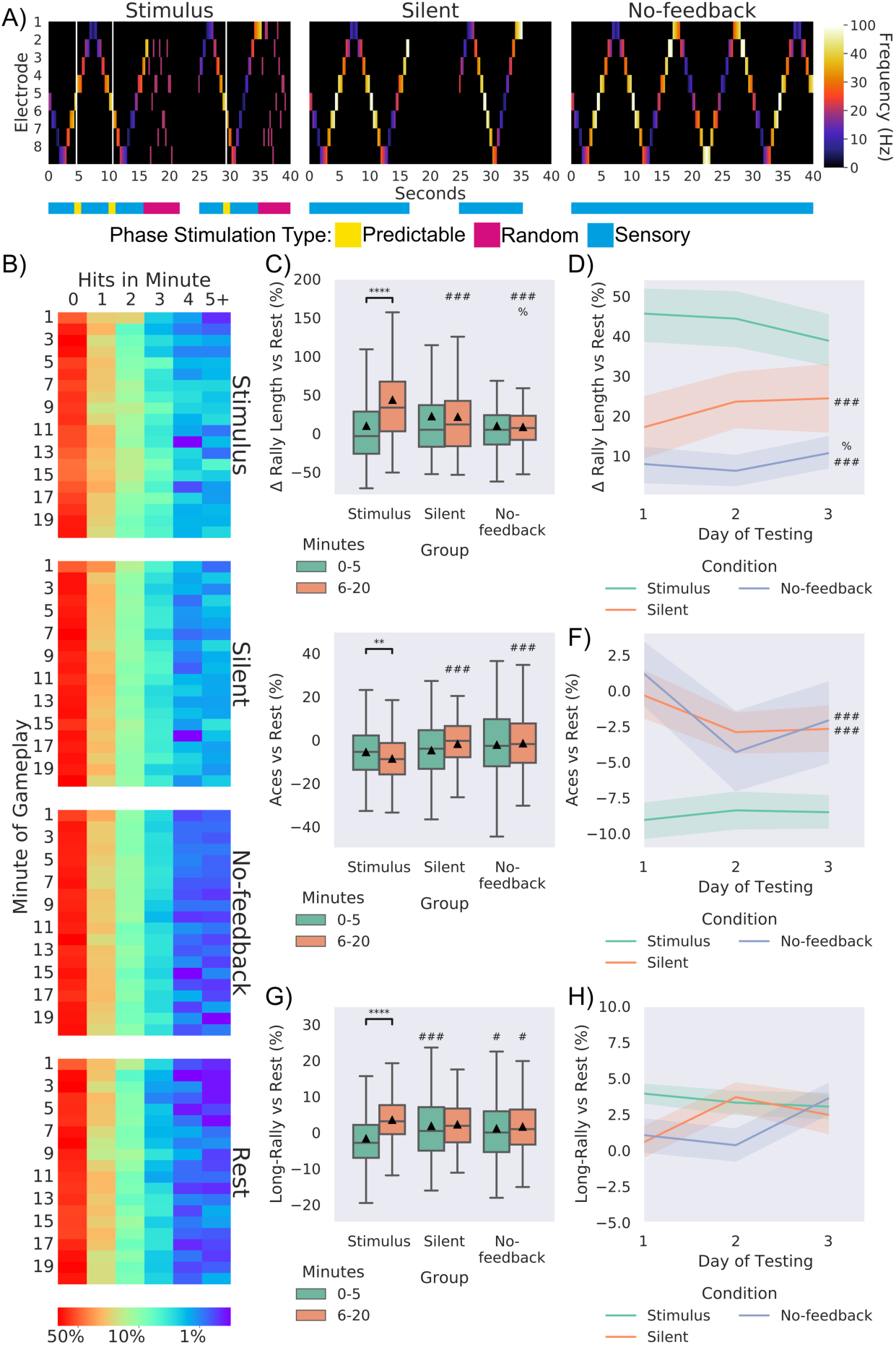
The Importance of Feedback in Learning. Significance bars show within group differences denoted with *. Symbols show between group differences at the given timepoint: # = *vs* Stimulus; % = *vs* Silent. The number of symbols denotes the p-value cut off, where 1 = p < 0.05, 2 = p <0.01, 3 = p < 0.001 and 4 = p <0.0001. Box plots show interquartile range, with bars demonstrating 1.5X interquartile range, the line marks the median and ▴ marks the mean. Errors bands = 1 SE. **A)** Schematic showing the stimulation from the 8 sensory electrodes across 40 seconds of the same gameplay for each of the three conditions. The bar below colour codes what phase of stimulation is being delivered. Where random stimulation follows a miss and predictable stimulation follows a hit in the Stimulus condition. Note the corresponding absence of any stimulation in the Silent condition and the of any change in sensory stimulation in the No-feedback condition. **B)** displays the probability of a certain number of hits occurring in a group at a specific minute. **C)** Using different feedback schedules the stimulus feedback condition showed significant learning (as in Figure 5A; *t* = 7.48, p = 1.58^-12^) and outperformed Silent and No-feedback average rally length, Silent feedback also showed higher performance compared to these groups at timepoint 2. **D)** displays this difference across day. **E)** Shows similar differences vs rest performance for aces across conditions, where the Stimulus group showed significantly fewer aces across time (*t* = 3.21, p = 0.002) **F)** displays this data across day. **G)** shows that the Stimulus condition showed significant increase (*t* = 3.21, p = 0.002) across timepoints, however as in **H)** no differences were found across time for long rallies.

Stimulus and Silent conditions showed overall higher performance compared to Rest and No-feedback conditions (Figure 7B). When testing for differences between groups in the percentage increase of average rally length over matched rest controls, a significant interaction was found (Figure 7C; Table S1). Only the Stimulus condition showed a significant increase in average rally length over time. While no differences were found for the first timepoint, a significant main effect of group was found at the second timepoint, where the Stimulus condition performed significantly higher than the Silent and No-feedback conditions (Table S2). Interestingly, the Silent condition also significantly outperformed the No-feedback conditions, although with less magnitude (Table S2). Importantly, this demonstrates that information alone is not sufficient; feedback is required to form a closed loop learning system. When followed up at the level of day for the second timepoint (Figure 7D) no significant differences over time were observed, but the between group differences were still observed. This trend was replicated when looking at aces both summed (Figure 7E) and across days of testing (Figure 7F). For long rallies the Stimulus group at timepoint 1 showed significantly fewer long-rallies compared to the Silent and No-feedback condition, being reversed at timepoint 2 with the Stimulus group showing significant more long rallies compared to the No-feedback condition (Figure 7G). No difference was found when this was followed up across day (Figure 7H). We also demonstrate that this learning is not seen in electrically inactive non-neural cells (Figure S4). Collectively this data establishes that adaptive behaviour seen in cortical cells altering activity to manipulate the environment can be an emergent property of engaging with—and implicitly modelling—the environment.

### Electrophysiological symmetry in latent activity is linked with higher performance

To determine whether spontaneous action potentials correlated with performance exploratory uncorrected Pearson’s correlations were computed for key activity metrics and average rally length in the last 15 minutes of gameplay. A significant positive correlation between mean firing and performance (Figure 8A) was found indicating a higher mean firing was associated with better performance, although max firing (Figure 8B) did not significantly correlate. This suggests that having well balanced higher activity was related to better performance, although the correlation was notably moderate. To further investigate whether the topographical distribution of activity correlated with performance, the absolute values of four discrete cosine transform (DCT) coefficients normalised to mean activity, was used to summarise spatial modes of spontaneous activity and assess symmetry of activity (Figure 8C). DCT (0,2) which shows difference between activity on the lateral edges and the lateral centre (Figure 8E) was significantly negatively correlated with performance. However, DCT (0,1) which measures activity across the horizontal plane (Figure 8C), DCT (1,0) which measures activity across the vertical plane (Figure 8F), and DCT (2,0) which measures activity on the horizontal edges vs the horizontal centre, did not significantly correlate. These correlations indicate that symmetrical activity across cultures underwrites better performance, but max activity does not. Given the distribution of the motor regions and sensory information, this finding is very coherent, as if there are no active cells in an area to either record signal from or deliver stimulation too, it would result in a dysfunctional system.

**Fig 8.**
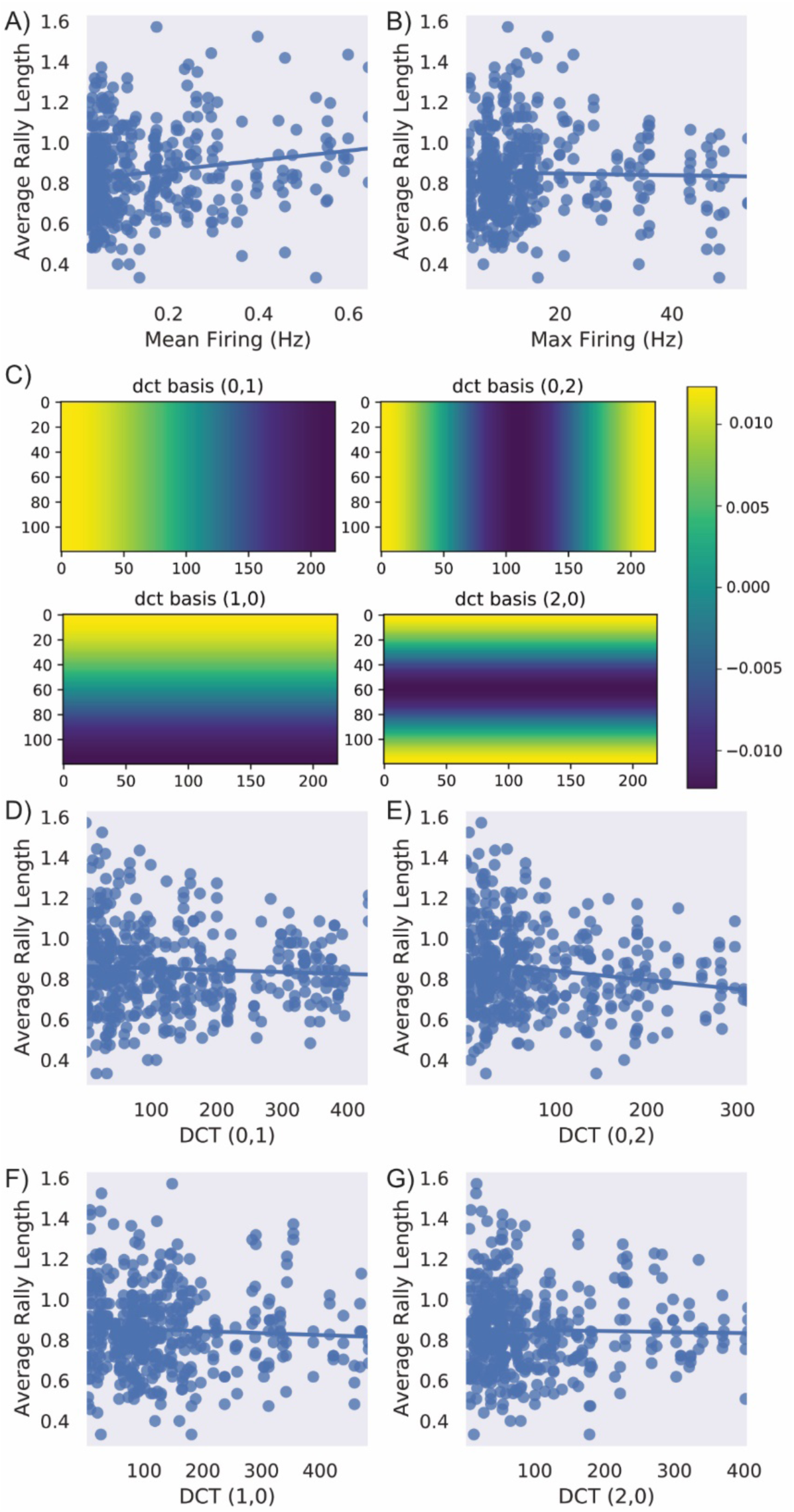
Relationship Between Latent Electrophysiological Activity for Higher Performance and the Importance of Feedback in Learning. **A)** A significant positive correlation between mean firing and performance was found (*r* = 0.17, *p* < 0.001) indicating a higher mean firing was associated with better performance. **B)** No significant relationship was found between max firing and performance. **C)** Absolute DCT values were calculated to determine whether there was a link between the layout of activity and performance. This shows how these DCT values were calculated for each type of score **D)** No significant relationship was found between DCT (0,1), **F)** DCT (1,0), or **G)** DCT (2,0). However, **E)** shows DCT 0,2 which mesures the difference between activity on the lateral edges and the lateral centre was significantly negatively correlated (*r* = -0.17, *p* < 0.001) with performance.

## DISCUSSION

Here we present a system, *Dishbrain*, which is capable of embodying neurons, from any source, in a virtual environment and measuring their responses to stimuli in real time. The ability of neurons, especially in assemblies, to respond to external stimuli in an adaptive manner is well established *in vivo*. However, this work is the first to establish this fundamental behaviour *in vitro*. We were able to use this silico-biological assay to investigate the fundamentals of neuronal computation. In brief, we demonstrate the first SBI device to show adaptive behaviour in real-time. The system itself offers opportunities to expand upon previous *in silico* models of neural behaviour, such as where models of hippocampal and entorhinal cells were tested in solving spatial and non-spatial problems (Sanders et al., 2020; Whittington et al., 2020). Minor variations on the *DishBrain* platform and selected cell type would enable an *in vitro* test to gain data around how cells process and compute information that was previously unattainable.

An example of this can be seen in the contrasting results between different cell sources. Active cortical cultures, from both human and mouse cell sources, both displayed synchronous activity patterns, in line with previous research^17–19,27^. Yet importantly, significant differences between cell sources were observed as human cortical cells always outperformed mouse cortical cells with nuances in gameplay characteristics. Although further work is required, this is the first work finding empirical evidence supporting the hypothesis that human neurons have superior information-processing capacity over rodent neurons^28,29^. This inherent difference between cell sources has been proposed to be due to denser and longer dendritic trees in human neurons, compared to mouse, which would yield different input-output properties and may thereby explain different computational capacities^30^. It was not previously possible to separate the neuroanatomical structure of different species from the microscopic structure of neurons in terms of computational power. Our work demonstrates that even when all key features are kept constant (cell number, sensory input, motor output etc.), there are key differences between human and rodent cortical neurons. This provides the first empirical evidence of differences in computational power between neurons from different species offering an exciting avenue for future research. Another finding from this work relates to innate cell organisation, seen in the definition of motor regions. Previously, motor regions were mapped from a population coding approach that incorporated spatial information following a network activity scan^5^. While our early pilot studies were similar, we focused on the extent that self-organisation would adapt if motor regions were fixed between cultures. When the EXP3 algorithm was used we found that experimental cultures showed significant preferences for layouts that could leverage biological processes like lateral inhibition. This is consistent with past work that finds feedback between environment and action is required for proper neural development^7^. However, it further suggests that perhaps this development occurs based on properties inherent at the level of the cell. This system provides the opportunity to explore network dynamics to better understand this aspect of self-organisation. At the technical level, this system is readily adaptable to include investigations into structural organisation of neural networks in both a physical and computational sense.

Most significantly, this work represents a substantial technical advancement in creating closed-loop environments for BNNs^5,6,31^. Here, we have emphasised the requirement for embodiment in neural systems for learning to occur. This is seen most significantly in the relative performance over experiments, where richer information and better feedback resulted in increased performance. Likewise, when no-feedback was provided yet information on ball position was available, cultures showed significantly poorer performance and no learning. Of particular interest was the finding that when stimulatory feedback was removed and replaced with silent feedback (i.e., the removal of all stimuli), cultures were still able to outperform those with no feedback as in the open-loop condition, albeit at a lesser extent. One interpretation is that playing ’pong’ generates more predictable outcomes than not playing ’pong’. Despite the outcome of a ‘failure’ not being unpredictable stimulation, given that the ball resets and the direction of the next movement is itself also unpredictable, this likely results in increased informational entropy, albeit to a far less extent. This is coherent with our results, as the more unpredictable an outcome, the greater the observed learning effect. However, the action of the BNN must have an outcome observable by the system. Therefore, it is coherent that the open-loop condition, which is by its nature the most predictable condition, did not result in learning. Stimulus alone is not sufficient to drive learning, there must be a motivation for the learning where altered behaviour can influence the future observable stimulus. When faced with unpredictable stimulus following unsuccessful performance, playing ’pong’ successfully acts as a free energy minimising solution. This offers a rather deflationary account of all goal-directed behaviour as the goal is just to minimise surprise. A key aspect of active inference is the selection of actions that minimise free energy expected following that action.

On this mechanistic level, we sought to demonstrate the utility of the *DishBrain* by testing base principles behind the idea of active inference via the FEP for intelligence, finding robust support for it. The closest previous work included studies of blind source separation in neural cultures^9,10^. However, this study did not offer physiologically plausible training environments and the system effectively existed in an open-loop environment. This makes any interpretation that the system in these studies was operating under the FEP difficult as changes in the external environment was not related back to the internal system of the neuronal cultures. Our work here demonstrates that when supplying unpredictable (random) sensory input following an ’undesirable’ outcome—and providing predictable input following a ’desirable’ one, we were able to significantly shape the behaviour of neural cultures in real time. The predictable stimulation could also be read as a stabilising synaptic weights in line with previous research^32,33^—or, in a complementary fashion, destabilising connectivity by destroying ’undesirable’ free energy minima. This may be a potential mechanism behind the FEP account of biological self-organisation, sometimes discussed in terms of *autovitiation* (i.e. self-organised instability by the destruction of self-induced but surprising fixed points of attraction)^8^. Crucially, expected free energy^13,34^ corresponds to uncertainty (i.e., informational entropy). This means that uncertainty minimising behaviour will have a natural curiosity, in the sense that it is necessarily information seeking. This is closely related to artificial curiosity in machine learning^35–37^ and intrinsic motivation in robotics^38^.

Due to current hardware limitations, the sensory stimulation is magnitudes coarser compared to that for *in vivo* organisms. Additionally, it was infeasible to meaningfully implement mechanisms that would be crucial for an *in vivo* organism attempting a comparable task, such as proprioception. Moreover, the relatively small number of cells embedded in a monolayer format means the neural architecture driving this behaviour is incredibly simple, in terms of the number of possible connections available compared to even small organisms that have a 3D brain structure. Nonetheless, using only simple patterns of predictable and unpredictable stimulation, this system was able to shape behaviour in an order of minutes. While within session learning was well established, between session learning over multiple days was not observed so robustly. Cultures appeared to relearn associations, with each new session. Given that cortical cells were selected, this is to be expected. *In vivo* cortical cells are not known to be specialised for long-term memory^39^. Future work with this system can investigate the use of other neuronal cell types and/or more complex biological structures.

## Conclusion

Using this *DishBrain* system, we have demonstrated that a single layer of *in vitro* cortical neurons can self-organise and display intelligent and sentient behaviour when embodied in a simulated game-world. We have shown that even without a substantial filtering of cellular activity, statistically robust differences over time and against controls could be observed in the behaviour of neuronal cultures in adapting to goal directed tasks. These findings provide a convincing demonstration of the SBI based system to learn over time in a goal-orientated manner directed by input. The system provides the capability for a fully visualised model of learning, where unique environments may be developed to assess the actual computations being performed by BNNs. This is something long sought after and extends beyond purely *in silico* models or predictions of molecular pathways alone (Karr et al., 2012; Whittington et al., 2020; Yu et al., 2018). Therefore, this work provides empirical evidence which can be used to support or challenge theories explaining how the brain interacts with the world and intelligence in general^11,40^. Ultimately, although substantial hardware, software, and wetware engineering is obviously still required to improve the *DishBrain* system, this work does evince the computational power of living neurons to learn adaptively in active exchange with their sensorium. This represents the largest step to date of achieving synthetic sentience capable of true generalised intelligence.

## METHODS

### Ethics statement

All experimental procedures were conducted in accordance with the Australian National Statement on Ethical Conduct in Human Research (2007) and the Australian Code for the Care and Use of Animals for scientific Purposes (2013) as required. Animal work was done under ethical approval E/1876/2019/M from the Alfred Research Alliance Animal Ethics Committee B. Experiments were performed at Monash University, Alfred Hospital Prescient with the appropriate personal and project licences and approvals. Work done using hiPSCs was done in keeping with the described material transfer agreement below.

### Experimental Procedures

No statistical methods were used to predetermine sample size. As all work was conducted within controlled environments uninfluenced by experimenter bias, experiments were not randomized, and investigators were not blinded to experimental condition. However, conditions were blinded before final analysis to prevent bias during analysis. Figure S5A presents a schematic of the overall experimental setup.

### Animal Breeding and maintenance

BL6/C57 mice were mated at Monash Animal Research Platform (MARP). Upon confirmation of pregnancy animals were transported via an approved carrier to the Alfred Medical Research and Education Precinct (AMREP). Pregnant animals were housed in individually ventilated cages until the date when they were humanely killed, and primary cells were harvested.

### Primary Cell Culturing

Cortical cells were disassociated from the cortices of E15.5 mouse embryos. Embryos were decapitated and with the aid of a stereotactic microscope the skin, bone and meninges were removed, and the anterior part of the cortex dissected out. Approximately 800,000 cells were plated down onto each pre-prepared HD-MEA. Cultures began to upregulate spontaneous activity and display synchronised firing around DIV 10 at which point they were used for experimentation.

### Stem Cell Lines

Initial work was conducted using a control hiPSC line supplied by the Gene Editing Facility at the Murdoch Children’s Research Institute (ATCC^®^ PCS-201-010) from an ATCC PCS-201-010 background and transferred under a Material Transfer Agreement. Later work involved an hiPSC lines used in this work constitutively expressing fluorescent reporters under control of the glyceraldehyde 3-phosphate dehydrogenase (GAPDH) promoter (cell lines were generated by Professor Edouard G. Stanley and colleagues from the Murdoch Children’s Research Institute and provided under a Material Transfer Agreement)^41^. The GAPDH gene encodes a protein critical in the glycolytic pathway, whereby ATP is synthesised from glucose. As this function is highly conserved across multiple cell types GAPDH is ubiquitously expressed at high levels across multiple cell types, making it a suitable gene for which to base a gene-expression system ^42^. This transgene expression system, termed GAPTrap, involves the insertion of the specific reporter gene into the GAPDH locus in hiPSCs using gene-editing technology^41^. For this study, RM3.5 GT-GFP-01 constitutively expressing green fluorescent protein under the GAPDH promoter was utilised. The RM3.5 hiPSC line was initially derived from human foreskin fibroblasts and reprogrammed using the hSTEMCCAloxP four factor lentiviral vector as reported previously^43^. All procedures described below were applied to be both cell lines. Both lines were maintained in an undifferentiated, pluripotent state in a feeder-free system using E8 media (StemCell Technologies, Canada) supplemented by a Penicillin/streptomycin solution at 5 µl/ml. Cells were plated on T25 353108 Blue Vented Falcon Flasks (Corning, Durham, USA) that were coated approximately 1 hr prior with the extracellular matrix vitronectin (Thermo Fisher Scientific, Carlsbad, USA).

### Stem Cell Maintenance

All procedures were carried out using sterile techniques. Prior to passaging cell confluence was recorded and the required split ratio was determined. Media was aspirated from cells and cells were washed with 5 ml of PBS -/- before passaging to remove detached cells and other debris. 3 ml of a 0.05 µM EDTA in PBS -/- was used for the dissociation and passaging of hiPSCs as aggregates without manual selection or scraping, was added to cells, and allowed to incubate at 37°C for approximately 3.5 mins. After visual examination using 10X microscope indicated that cells had lost sufficient adhesion, EDTA was aspirated, and blunt trauma applied to base of the T25 flask to dislodge cells. Cells were suspended in 2 ml E8 and transferred to 15 ml falcon tube. As described above, vitronectin coated T25 flasks were prepared and aspirated before the addition of 5 ml of E8 solution. Approximately 1:10 of evenly distributed cell suspension was added to the prepared T25 flask. The flask was then gently swirled to ensure even distribution before being incubated overnight at 37°C. Media was changed daily.

### Stem Cell Dual SMAD Differentiation

Cellular differentiation followed a titrated dual SMAD inhibition protocol for the generation of cortical cells from pluripotent cells established by the Livesey group with minor adjustments as represented in Figure S5B^19^. Cells were plated in 24 well plates coated with human laminin H521. When cells reached ≈80% confluency, neural induction was initiated by using standard neural maintenance (N2B27) Base Media with 100 ng/ml LDN193189 (Stemcell Technologies Australia, Melbourne, Australia) and 10 µm SB431542 (Stemcell Technologies Australia, Melbourne, Australia). Media was changed every day from day 0 to day 12. After appearance of neural rosettes and initial passaging standard N2B27 media with FGF2 20 ng/ml was utilised from day 12 to day 17 to achieve a dorsal forebrain patterning. Cells were then expanded and deemed ready for plating onto MEA or slides based on morphology at approximately 30 – 33 days. On the day of transplant, cells were detached with Accutase (Stemcell Technologies Australia, Melbourne, Australia) to a single cell suspension and centrifuged at 300*g*. The cell pellet was resuspended at 10,000 cells/µl in BrainPhys (Stemcell Technologies Australia, Melbourne, Australia) neural maintenance media with Rho Kinase Inhibitor IV (Stemcell Technologies Australia, Melbourne, Australia;1:50 dilution) with approximately 10^6^ cells plated onto each MEA. Cells began to display early but widespread spontaneous activity around DIV 80, at which point they were ready for experimentation.

### Stem Cell NGN2 Direct Differentiation

Cortical excitatory neurons were generated by the expression of NGN2 in iPSCs. IPSCs were plated at 25,000 cells/cm^2^ in a 24-well plate coated with 15µg/ml human laminin (Sigma, USA). The following day, cells were transduced with NGN2 lentivirus (containing a tetracycline-controlled promoter coupled with a puromycin selection cassette) in combination with a lentivirus for the rtTA (reverse tetracycline-controlled transactivator). NGN2 gene expression was activated by the addition of 1 µg/ml doxycycline (Sigma, Australia), this was referred to as differentiation day 0. Cells were cultured in neural media consisting of 1:1 ratio of DMEM/F12:Neurobasal media supplemented with (all reagents from Thermofisher, USA) B27 (#17504-044), N2 (17502-048), Glutamax (#35050-060), NEAA (#11140-050), β-mercaptoethanol, ITS-A (#51300-044) and penicillin/streptomycin (#15140-122). On Day 1 1.0µg/ml puromycin (Sigma, Australia) was added for 3 days at which point neurons were supplemented with 10µg/ml BDNF (Peprotech, USA) and lifted with accutase, in preparation for plating on HD-MEA chips. HD-MEA chips were pre-treated with 100µg/ml PDL (Sigma, USA) and 15µg/ml laminin (Sigma, USA). For each well 1×10^5^ NGN2 induced neurons at DD4 were combined with 2.5×10^4^ primary human astrocytes (ScienceCell, USA) in each well of the MEA plate. To arrest cell division of astrocytes 2.5µM Ara-C hydrochloride (Sigma, USA) was added at day 5 for 48 hours. Cells were maintained in neural media supplemented with BDNF and media changed at least 1 day prior to recordings.

### MEA setup and preparation

MaxOne Multielectrode Arrays (MEA; Maxwell Biosystems, AG, Switzerland) were used for this research. The MaxOne is a high-resolution electrophysiology platform featuring 26,000 platinum electrodes arranged over an 8 mm^2^. The MaxOne system is based on complementary meta-oxide-semiconductor (CMOS) technology and allows recording from up to 1024 channels and stimulation from up to 32 units. MEAs and chambered glass slides are coated with either polyethylenimine (PEI) in borate buffer for primary culture cells or Poly-D-Lysine for cells from an iPSC background before being coated with either 10 µg/ml mouse laminin or 5 µg/ml human 521 Laminin (Stemcell Technologies Australia, Melbourne, Australia) respectively to facilitate cell adhesion.

### Plating and Maintaining Cells on MEA

Approximately 10^6^ cells were plated on MEA after preparation via method already described. Cells were allowed approximately one hour to adhere to MEA surface before the well was flooded. The day after plating, cell culture media was changed to BrainPhys™ Neuronal Medium (Stemcell Technologies Australia, Melbourne, Australia) supplemented with 1% penicillin-streptomycin. Cultures were maintained in a low O_2_ incubator kept at 5% CO_2_, 5% O_2_, 36°C and 80% relative humidity. Every two days, half the media from each well was removed and replaced with free media. Media changes always occurred after all recording sessions.

### Measuring of Electrophysiological Activity

Licenced MaxLab Live Scope V20.1 software was used to run activity scans. Checkerboard assays consisting of 14 configurations at 15 seconds of spike only record time were run daily immediately preceding the running of the DishBrain software. Gain was set to 512x with a 300 Hz high pass filter. Spike threshold was set to be a signal six sigma greater than background noise as per recommended software settings. Mean, max and variance of both amplitudes and firing rates was extracted from these assays and mapped using custom software: the first nine components of discrete cosine transform basis functions of space were used to summarise the spatial profile of spiking activity. The ensuing coefficients were then used in subsequent correlation analyses.

### *DishBrain* software platform

The current *DishBrain* platforms is configured as a low-latency, real-time MEA control system with on-line spike detection and recording software. See Figure S3 and **Supplementary Text 1**. The DishBrain software runs at 20,000 Hz and allows recording at this incredibly fine timescale. Working closely with MaxWell Biosystems we enabled capabilities not available using the native vendor software. The existing API was used only for loading configurations. Low level code was written in C to allow for minimal processing latencies—so that packet processing latency was typically <50 µs. High level code, including configuration set ups and broader instructions for game settings were implemented in Python. **Figure S5C** shows an image of the game visualiser, and a real-time interactive version is available at https://spikestream.corticallabs.com/. This allowed a spike-to-stim latency of approximately 5 ms, with the substantive delay due to inflexible hardware buffering built into MaxOne hardware. Where appropriate, the EXP3 machine learning algorithm was used to sample two predefined motor regions to select the best configuration to interpret movement commands for the paddle. When the system failed to move the ‘paddle’ into a correct position (to contact the ball), a random stimulus was applied to culture at 5 Hz and 150 mV. After a 1 s delay—to allow the culture to recover—play was resumed. The online spike detection software was developed using an adaptive threshold-based detector. The threshold was typically set at 6 sigmas above noise estimates. We established this use a mean absolute deviation (MAD) estimate, which was multiplied by a correction factor. Infinite Impulse Response (IIR) filters and Digital Signal Processing (DSP) were used to prepare raw signals for spike detection.

### Input configuration

Stimulation is delivered at a given Hz and voltage as described in the main text to key electrodes in a sensory area, as shown in Figure 4B. Initial experiments delivered purely place-coded stimulation, where the distance from the centre of the sensory area was interpreted as distance from the centre of the paddle aligning with the ball. As described in the main text, later experiments adopted a mixed coding scheme, where the place coding was combined with a rate coding that delivered stimuli at 4 Hz when the ball was closest to the opposing wall and increased to a max of 40 Hz as the ball reached the paddle wall.

### Output configuration

Initially two predefined motor regions were defined on the MEA. Activity was measured over these two regions, where the region with higher activity would move the paddle in a corresponding direction. This was found to be extremely sensitive to culture characteristics, where asymmetrical spontaneous spiking activity in cultures would cause the paddle to move swiftly in only one direction. To counter this, a gain function was implemented, which measured activity in both regions and added a multiplier to a target of 20 Hz. Activity >20 Hz was weighted by a correction factor >1, while activity <20 Hz was weighted by a correction factor <1. An EXP3 algorithm was implemented to select the different configuration options illustrated in Figure S3^44^. We found that experimental cultures preferred configuration 3, while media only control cultures preferred configuration 0. As such, configuration 3 was selected as it offered the possibility for biologically relevant features, such as lateral inhibition, and minimised the chance of apparently successful performance through bias alone—as it precludes a direct relationship between input stimulation and output activity recording.

### EXP3 Algorithm

An Exponential-weight algorithm for Exploration and Exploitation (EXP3) algorithm was used initially for the adaptive selection of electrode layouts, with the objective of optimising gameplay performance ^45^. This algorithm was implemented to maintain a list of weights for each action and was designed to minimise regret by preferencing electrode configurations which were associated with a higher probability of the ball being returned. This is described in detail in **Supplementary Text 2**.

### Immunocytochemistry

Cells were washed three times with sterile PBS and then fixed using 4% PFA for 20 mins. After washing cells were blocked 0.3% Triton-X and 1% goat serum in PBS for 1 hr. Primary antibodies specific for Synapsin1 (1:500; ab254349; Rabbit; Abcam, Cambridge, MA, USA), NeuN (1:500; ab104225; Rabbit; Abcam, Cambridge, MA, USA), Beta-III Tubulin (1:500; MAB1637, Mouse; Kenilworth, NJ, USA), MAP2 (1:1000; Chicken; ab5392; Abcam, Cambridge, MA, USA), TBR1 (1:200; ab183032; Rabbit; Abcam, Cambridge, MA, USA), GFAP (1:500; ab4674; Chicken; Abcam, Cambridge, MA, USA), and KI67 (1:500; ab245113; Mouse; Abcam, Cambridge, MA, USA) were applied overnight. After washing, secondary antibodies (chicken 555, rabbit 488, mouse 647; Abcam, Cambridge, MA, USA) were incubated for 2 hrs. This was followed by 10 mins of DAPI Staining Solution in PBS (1:1000, ab228549, Abcam, Cambridge, MA, USA) after which point slides were cover-slipped with ProLong Gold Antifade Mountant (Thermo Fisher Scientific, Waltham, MA, USA) mounting media and allowed to dry for 48 hrs. **Scanning Electron Microscopy.** At various designated endpoints, media was aspirated from the MEA wells and cells were fixed with 2.5% glutaraldehyde (Electron Microscopy Sciences, PA, USA) and 2% paraformaldehyde (Electron Microscopy Sciences, PA, USA) in a 1 M sodium cacodylate buffer for 1 hr. They were then washed three times in 1 M sodium cacodylate buffer before being post-fixed with 1% OsO_4_ in a 1M sodium cacodylate buffer for 1 hr. OsO_4_ was removed and the fixed cells were washed with three times in milliQ water and dehydrated via an ethanol gradient exchange (30%, 50%, 70%, 90%, 100%, 100% v/v) for 15 mins each. After dehydration, the cells were dried by hexamethyldisilazane (Sigma Aldrich, St. Louis, MO, USA) exchange (3×10 mins), and then allowed to evaporate for 5-10 mins. MEA chips were then affixed to an aluminium stub with carbon tape and sputter coated with 30 nm layer of gold using a BAL-TEC SCD-005 gold sputter coated. All procedures were performed at room temperature. Coated MEA chips were then imaged using a FEI Nova NanoSEM 450 FEGSEM operating with an acceleration voltage of 10 kV and a working distance of 12 mm. Images were then analysed using ImageJ v. 1.52k and false coloured using Adobe Photoshop.

### Widefield fluorescence microscopy

Images were captured using a Nikon Ti-E upright light microscope equipped with a motorised stage. All widefield images were captured using a 20X objective.

### Data analysis

Data was analysed using custom code written in Python. Error bars are described in captions, except where graphs are box and whisker plots, where the line is the median, box indicates lower quartile to upper quartile and error bars show the rest of the distribution excluding outliers. The illustrative data provided in the text and figures include means and standard deviations. An alpha of *p* < 0.05 was adopted to establish statistical significance, providing a 5% chance of a false positive error. Where suitable assumptions were met, inferential frequentist statistics were used to determine whether statistically significant differences existed between groups. All tests were two tailed tests for statistical significance. For related samples *t*-tests or independent *T*-tests alpha values for significance were corrected via the Bonferroni method. For one-way analysis of variance (ANOVA) and the multivariate 2 x 3 repeated measures ANOVA when a significant interaction or main effect was found, this was followed up with pairwise Games-Howell post hoc tests with Tukey correction for multiple comparisons. This was adopted as = there were always differences between sample sizes and variance due to inclusion of in-silico controls. As seen in **Figure S5D**, four DCT basis functions were used to summarise spatial modes of spontaneous activity. Pairwise Pearson’s correlations were used to test the relationship between the ensuing scores—along with time (s) and max and mean firing rates (Hz)—with average rally length.

## Acknowledgements

The authors wish to acknowledge and thank Professor Anthony N. Burkitt, Professor David Walker, Professor Adeel Razi, Dr Chris French and Dr Alberto Roselló-Díez for their advice and comments on the manuscript. The authors acknowledge and thank Professor Edouard G. Stanley and Professor Andrew Elefanty from the Murdoch Children’s Research Institute (MCRI) for their provision of RM3.5 cells along with Dr Ana Antonic-Baker for their assistance. The authors acknowledge the use of instruments and assistance at the Monash Ramaciotti Centre for Cryo-Electron Microscopy, a Node of Microscopy Australia. The authors acknowledge the use of instruments and Monash Micro Imaging (MMI) Facility and the associated assistance of Dr Stephen H. Cody and Dr Chad Johnson.

## Data Access

All data not considered proprietary material are available to share upon reasonable request to the corresponding author to assist with reproducibility.

## Custom Code Access

All custom code not considered proprietary material are available to share upon reasonable request to the corresponding author to assist with reproducibility.

## Author Contributions

Conceptualization, B.J.K., A.K., A.R. K.J.F; methodology, B.J.K., A.K.; software: A.K., B.J.K.; validation, B.J.K., N.T.T., B.J.P.; formal analysis, B.J.K.; cell culture, B.J.K., N.T.T., B.J.P., B.R.; investigation, B.J.K., N.T.T., B.J.P; data curation, B.J.K, A.K.; writing—original draft preparation, B.J.K; writing—review and editing, B.J.K., K.J.F., A.R., A.B., N.T.T., B.J.P., B.R.; visualization, B.J.K., N.T.T., B.J.P., B.R.; project administration, B.J.K.; supervision, B.J.K.

## Supplementary Text

### Supplementary Text 1: Development of the modular, real-time platform *DishBrain* to harness neuronal computation

The MaxOne MEA is not only capable of measuring changes in electrical activity brought about from action potentials, but also of stimulating cells at a range of voltages, in a manner which is relatively non-invasive to cells, and effectively elicits action potentials or responses in a comparable manner to internal electrical stimulation^26^. With the appropriate coding scheme, external electrical stimulations can convey a range of information, providing the capacity to not only ‘read’ information from a neural culture, but also to ‘write’ data into one. We set out to build a system, named ‘*DishBrain’* (Fig, S2A), which would allow us to integrate these two principles into a closed loop, in the hope of allowing the neurons to achieve embodiment and agency in a virtual environment, with demonstrable learning effects. The DishBrain system is controlled by a low latency, real-time piece of software named ‘*DishServer’*, which replaces and extends a corresponding piece of MaxWell vendor software called ‘MXWServer’. DishServer is capable of receiving voltage readings from MaxOne vendor hardware, processing these readings, simulating a virtual environment, encoding the results as MaxOne electrode commands, and sending these commands back to the MaxOne hardware. When run on a computer with access to a MaxOne hardware setup with a live culture in place, the system acts as a closed loop that we can configure and record for analysis. The system is easily adaptable to other MEA hardware and virtual environments as well, which could exhibit different learning and embodiment effects if tested.

So far, the main use of *DishBrain* has been to embody neural cultures in a simulation of the classic arcade game ‘pong’, with neuron activity read from multiple ‘motor regions’ defined by distinct subsets of the MEA, and sensory information encoded as stimulation at any of eight distinct stimulation sites placed opposite from those motor regions. The MaxOne MEA is configured to read up to a particular 1024 of its 26,400 electrodes, at a rate of 20,000 samples per second. As shown in Figure S2B, these samples are optionally recorded as-is, for later analysis, but are also run through a sequence of computationally efficient Infinite Impulse Response (IIR) filters to calculate noise and activity levels, which are compared in order to detect spikes. Incoming samples are filtered with a 2nd order high-pass Bessel filter with 100Hz cut-off, the absolute value is then smoothed using a 1st order low-pass Bessel filter with 1Hz cut-off, the spike threshold is proportional to this smoothed absolute value. Spikes are themselves optionally recorded, and either way are counted over a period of 10 milliseconds, (200 samples,) at which point the game environment is given the number of spikes detected in each of the configured electrodes, and these spike counts are interpreted as motor activity depending on which motor region the spikes occurred in, moving the paddle up or down. At each of these 10ms intervals the pong game is also updated, with a ball moving around a play area at a fixed speed, ‘bouncing’ off the edges of the play area and off the paddle, until it hits the edge of the play area behind the paddle, which marks the end of one ‘rally’ of pong. During each rally the location of the ball relative to the paddle is encoded as stimulation to one of eight stimulation sites, which is tracked in an internal ‘stimulation sequencer’ module. The stimulation sequencer is updated 20,000 times a second, once every time a sample is received from the MEA, and once the previous lot of MEA commands should have finished, it constructs another sequence of MEA commands based on the place-code and rate-code information that it has been configured to transmit. The stimulations take the form of a short square bi-phasic pulse that is a positive voltage, then a negative voltage. A Digital to Analog Converter (or DAC) on the MEA will read and apply this pulse sequence to the given electrode. To interface with Maxwell API, *DishBrain* use a negative DAC value first because this corresponds to a positive voltage in the MaxWell API. At the end of the rally, the game environment will instead configure the stimulation sequencer to apply stimulation at random sites, for a period of four seconds, followed by a configurable rest period of up to four seconds, followed then by the next rally. Finally, the spike detection is also capable of ‘blinding’, which is expected to occur after each stimulation; in order to prevent DAC stimulation from being interpreted as neuron activity, all 1024 channels are ignored for a configurable number of samples, after either detecting anomalous activity directly, or after receiving acknowledgement from the MEA that a DAC command has been executed.

Given the multitude of possible variations inherent in a system like this, it was necessary to fix some parameters and empirically test others. Stimulation is delivered at specific locations, frequency, and voltage to key electrodes in a topographically consistent manner in the sensory area relative to the current position of the paddle (Figure S3: Configuration 0). This was designed to mimic retinotopic and topographic representations commonly found in nearly all neural systems for representing the external world^46,47^. Other parameters, such as voltage were determined through empirical testing. Initial tests were conducted to assay which conditions cell cultures would survive. Testing time was found to be a highly sensitive parameter, as cells did not tolerate testing times >1.5 hours. When measurements were taken it was concluded that this was likely due to increased temperature in the well in which cells were plated in due to activity and the resulting increased evaporation and changes in osmolarity. To the surprise of the researchers, cells survived testing administration of stimulation up to 3000 mV for up to one hour which was the maximum testing time considered given the above findings. While this did create excess noise in recording cellular activity across the MEA during the stimulation period, there were no significant changes to spontaneous activity in the cell cultures before and after the period of stimuli administration. Given that cells appeared robust to voltage stimulation, the decision was made to base voltage levels on existing evidence of neurological function. To prevent forcing hyperpolarised cells from firing, 75 mV at 4 Hz was chosen as the sensory stimulation voltage that would relate to where the ball was relative to the paddle. In order to add unpredictable external stimulus into the system, when the culture fails to line the paddle up to connect with the ball, the ‘punishing’ stimulus was set at 150 mV voltage and 5 Hz. It was hypothesised that this higher voltage would be sufficient to force action potentials in cells subjected to the stimulation regardless of the state the cell was in, thereby being even more disruptive to the culture. Initially, two distinct areas were defined as ‘motor regions’, where activity in motor region 1 moved the paddle ‘left’ and activity in motor region 2 moved the paddle ‘right’. Due to the technical difficulty of culturing neurons that displayed perfectly symmetrical activity in both these regions it was found to be necessary to add ‘gain’ into our system. These took a real-time value based on the mean firing in each motor region and multiplied it to achieve a target value of 20 Hz across the entire region. This would allow changes in activity in each given region to influence the paddle position, even if they displayed different latent spontaneous activity.

### Supplementary Text 2: The development of an EXP3 algorithm to assess different motor region recording layouts and further refinement of blinding protocols

After initial pilot testing of the *DishBrain* system, two pathways were identified to modify performance: encoding of information and decoding of activity. The initial focus was to improve the latter. It was hypothesized that the simplified decoding system of measuring activity in two motor regions that were congruent where activity was stimulated might not only be inefficient but also prone to bias. To investigate this further an EXP3 machine learning algorithm was used to sample two predefined motor regions to select the best configuration from six possible configurations and interpret movement commands for this paddle. The EXP3 algorithm was used for adaptive selection of electrode layouts, with the objective of increasing the expected number of times that a culture is able to hit the ball in each pong rally. EXP3 is robust to changes in the underlying distribution of returns; this is important because neurons are also concurrently learning, and their behaviour changing over time. Optimising over all possible assignments of electrodes to actions would require a prohibitively large set of choices, so a representative set of balanced layouts were used. EXP3 is an online optimisation algorithm for the “multi-armed bandit” problem. It selects between several discrete choices, over a series of rounds. Each discrete choice yields an observable stochastic loss. The best choice is never revealed, even post-hoc. Quality of choices can only be inferred from noisy returns; exploration and exploitation must be balanced. In this work, one of a discrete set of electrode-action mappings called ’motor layouts’ was chosen on each round. The loss to be minimised is calculated using the following equation:

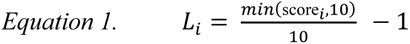

Where loss_i is the loss at the end of the rally i and score_i is number of bounces during that rally. During the i-th rally, layout_i is used and is fixed during the entire rally. At the end of the rally, layout_i+1 is chosen by EXP3 for the next rally and the game play continues. When using EXP3 the system can adaptively optimise performance by choosing from a fixed set of alternative motor layouts (Figure S3). At the same time, a new blinding method (consensus blind) based on blinding all signals when >15 simultaneous large (>75 mV) spikes were detected, was implemented to block stimulation delivered by the system from being registered as cellular activity. It was hypothesised that a lack of blinding administered signals may contribute to the apparent performance observed in controls in our pilot study. As described in the main text and shown in Table S3, experimental chips with configurations that would enable lateral inhibition were found to be selected significantly more compared to other configurations resulting in an equal distribution (*χ2* = 35690.93, *p*<0.0001), including those that were more simplified like that used in the pilot where activity on the left moved the paddle left and conversely for the right (Figure S3**: Configuration 0**) and would be most easily influenced by various sources of bias. This behaviour has been observed in experiments involving multiple human and mice neurological functions^48–50^. When the frequency tables of these two distributions were compared, they were also found to be significantly different, (*χ2* = 15229.323, *p*<0.0001). Considering these differences, it is not valid to compare experimental and control groups as they are operating off different types of configurations.

Given the apparent preference for configurations that would allow processes such as lateral inhibition to occur in experimental chips, coupled with the concern of having different groups operating from different configurations, it was decided to select configuration 3 for all cultures going forward, as it was chosen most frequently by the EXP3 algorithm. Moreover, if consensus blinding behaved as expected, control chips should also show no preference. This led us to suspect that consensus blinding was ineffective and on further investigation, particularly when using a higher and variable frequency of sensory stimulation we discovered more evidence of consensus blinding failing than our previous testing revealed. To counter this, a new blinding method was implemented, which was termed ‘command count blinding’. This method blinded our readout of all motor activity when a command was sent to generate any form of stimulation. During testing this was found to be significantly more robust than the previously used consensus blinding and allowed us to proceed with increasing the density and variability of sensory stimulation. A second method of gathering control data by gathering data during ‘rest’ periods was also implemented, whereby the game would continue with all events still being logged, but no stimulation was administered to the culture.

---

**Fig. S1.**
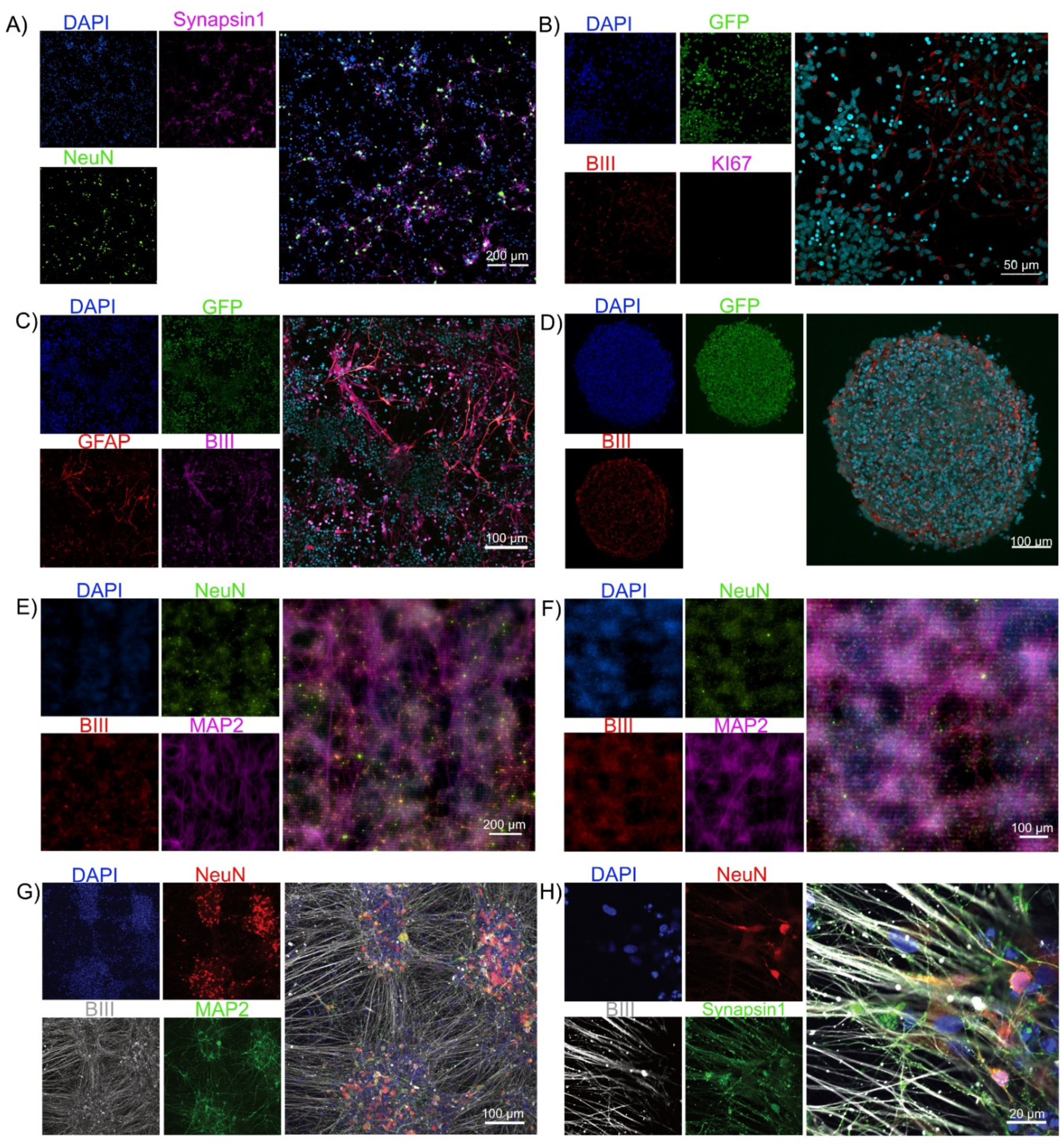
Cortical neurons can be obtained via multiple methods. Scale bars as shown on figure. **A)** Primary mouse cortical neurons show diverse expression of synapsin1 which marks synaptic vesicles and actin filaments across long reaching neural networks. **B)** – **F)** Shows that using a RM3.5 cell line comparable cortical cultures can be generated using the dual SMAD inhibition protocol described in Methods. **B)** Shows endogenous expression of GFP, BIII marking axons and a lack of Ki67 suggesting no dividing cells, **C)** additional shows these cells expressing GFAP for supporting glial cells Further images in **D)** show a characteristic neurosphere structure neurons would often spontaneously form when plated at high density, a dense pseudo three-dimensional sphere with dense connections of neurons and axons throughout. **E)** & **F)** display hIPCSs differentiated to neurons using the NGN2 method and mouse primary cortical neurons respectively, both plated of HD-MEA and allowed to mature before staining. These cells display all markers previously described, but due to the reflective material of the CMOS chip, it is infeasible to get high resolution fluorescent images of cells on the chips, leading to the adoption of SEM imaging shown in the main text. **G)** & **H)** also show hIPCSs differentiated to neurons using the NGN2 method; **G)** Staining of mature neural monolayer cultures with the majority of cells expressing NeuN which marks neuronal cells, MAP2 marks dendrites and β3-Tubulin which marks long-range axons. **H)** Further staining shows that along with β3-Tubulin these cells express the pre-synaptic marker synapsin1 across the soma and cell projections.

**Fig. S2.**
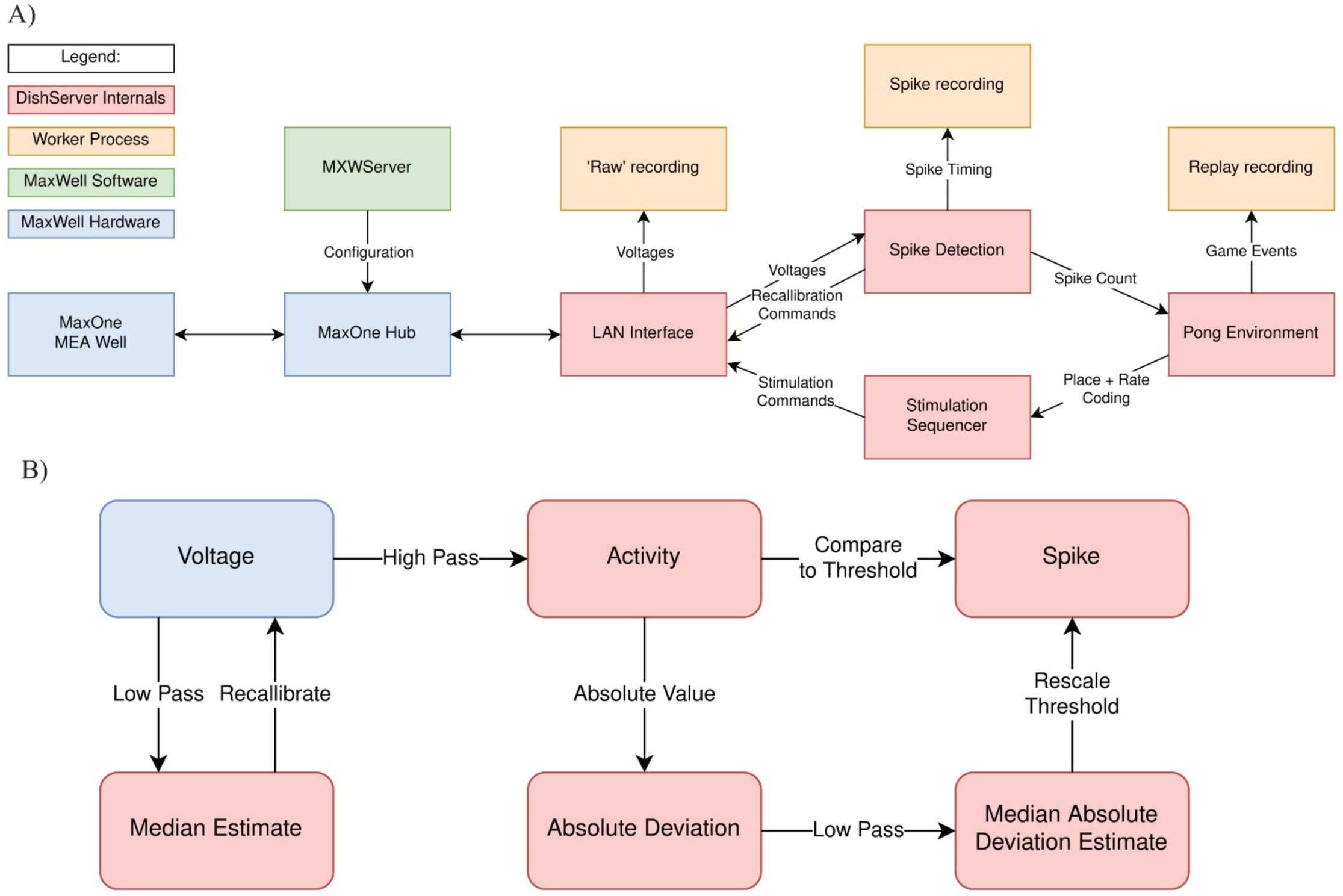
Schematics of software used for *DishBrain*. **A)** Software components and data flow in the DishBrain closed loop system. Voltage samples flow from the MEA to the ‘pong’ environment, and sensory information flows from the ‘pong’ environment back to the MEA, forming a closed loop. The blue rectangles mark proprietary pieces of hardware from MaxWell, including the MEA well which may contain a live culture of neurons. The green MXWServer is a piece of software provided by MaxWell which is used to configure the MEA and Hub, using a private API directly over the network. The red rectangles mark components of the ’DishServer’ program, a high-performance program consisting of four components designed to run asynchronously, despite being run on a single CPU thread. The ’LAN Interface’ component stores network state, for talking to the Hub, and produces arrays of voltage values for processing. Voltage values are passed to the ’Spike Detection’ component, which stores feedback values and spike counts, and passes recalibration commands back to the LAN Interface. When the pong environment is ready to run, it updates the state of the paddle based on the spike counts, updates the state of the ball based on its velocity and collision conditions, and reconfigures the stimulation sequencer based on the relative position of the ball and current state of the game. The stimulation sequencer stores and updates indices and countdowns relating to the stimulations it must produce and converts these into commands each time the corresponding countdown reaches zero, which are finally passed back to the LAN Interface, to send to the MEA system, closing the loop. The procedures associated with each component are run one after the other in a simple loop control flow, but the ‘pong’ environment only moves forward every 200^th^ update, short-circuiting otherwise. Additionally, up to three worker processes are launched in parallel, depending on which parts of the system need to be recorded. They receive data from the main thread via shared memory and write it to file, allowing the main thread to continue processing data without having to hand control to the operating system and back again. **B)** Numeric operations in the real-time spike detection component of the *DishBrain* closed loop system, including multiple IIR filters. Running a virtual environment in a closed loop imposes strict performance requirements, and digital signal processing is the main bottleneck of this system, with close to 40 MiB of data to process every second. Simple sequences of IIR digital filters is applied to incoming data, storing multiple arrays of 1024 feedback values in between each sample. First, spikes on the incoming data are detected by applying a high pass filter to determine the deviation of the activity, and comparing that to the MAD, which is itself calculated with a subsequent low pass filter. Then, a low pass filter is applied to the original data to determine whether the MEA hardware needs to be recalibrated, affecting future samples. This system was able to keep up with the incoming data on a single thread of an Intel Core i7-8809G.

**Fig. S3.**
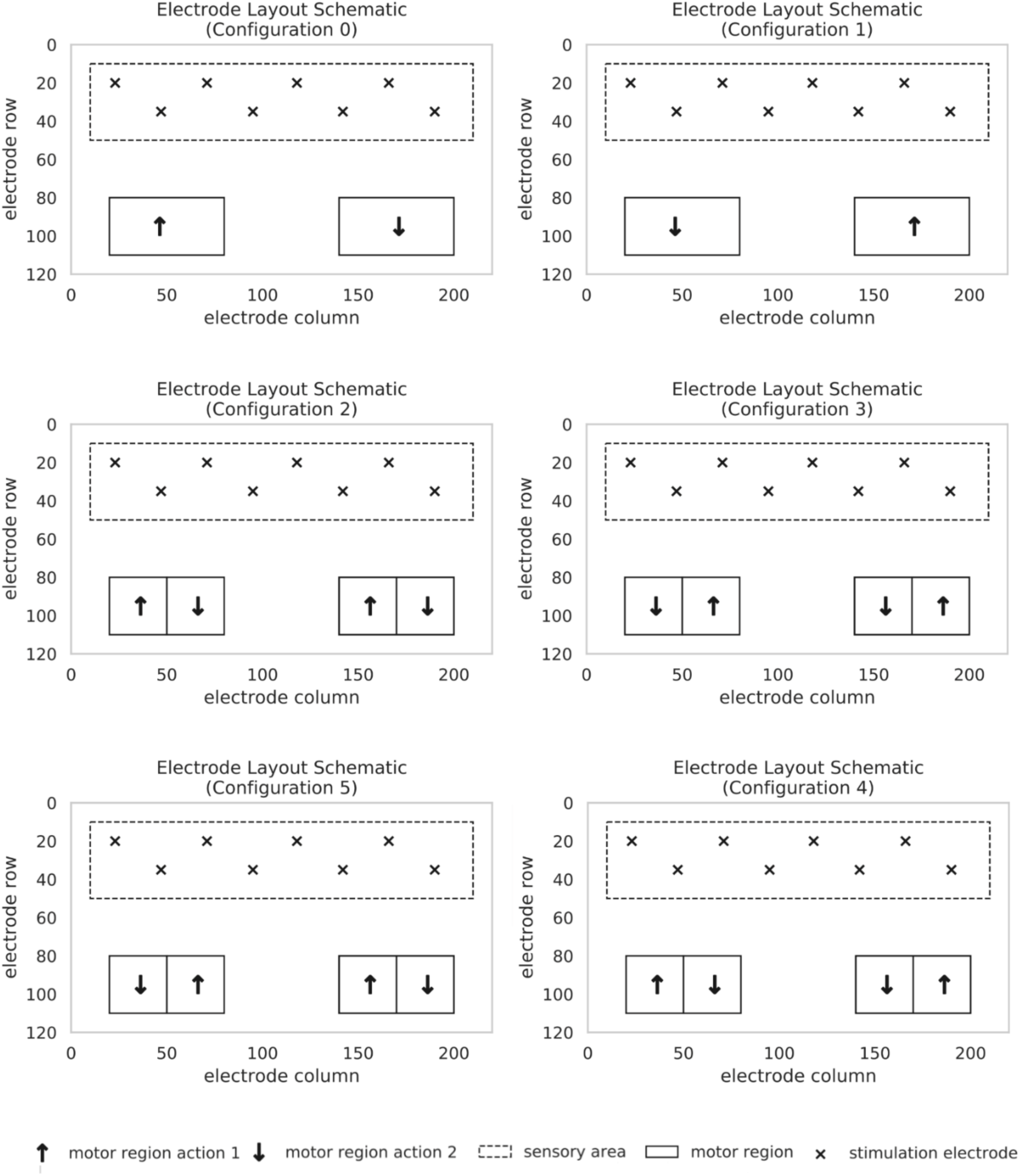
Representation of the specific configurations of the DishBrain platform. Stimulation is delivered to a predefined sensory area and activity is measured in the motor regions to determine how the paddle will move. Feedback is provided via the sensory area based on the outcome of the motor region activity. Note the different configurations in which motor activity may have been interpreted. Configuration 0 was initially adopted as the beginning choice, however when the EXP3 algorithm was used to control selection from all of the above options, experimental cultures adopted a preference for configuration 3, which was then adopted going forward.

**Fig S4.**
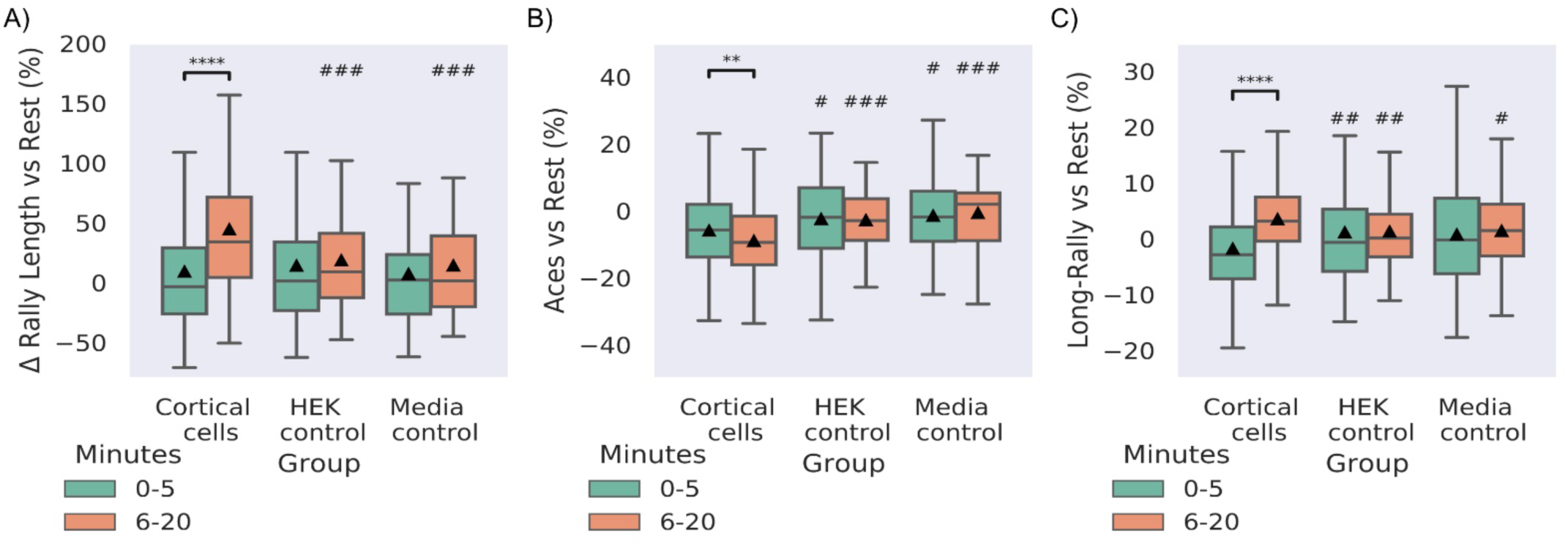
Electrically inactive non-neural cells also display no learning over time and perform at media control levels compared to cortical cells. Significance bars show within group differences denoted with *. Symbols show between group differences at the given timepoint: # = vs Cortical cells. The number of symbols denotes the p-value cut off, where 1 = p < 0.05, 2 = p <0.01, 3 = p < 0.001 and 4 = p <0.0001. Box plots show interquartile range, with bars demonstrating 1.5X interquartile range, the line marks the median and ▴marks the mean. **A)** Looking at the % change in rally length compared to match rest controls, cortical cells condition showed significant *t* = 8.22, p = 1.15^-15^) and outperformed HEK293T cells and media control groups at timepoint 2 which showed no change over time (Table S2). **B)** Shows similar differences vs rest performance for aces across conditions, where the Cortical cell group showed significantly less % of aces across time (*t* = 3.21, p = 0.002) along with significantly fewer aces than the HEK control and Media control groups at both timepoints (Table S2). **C)** differences vs rest performance for % if long-rallies across conditions, where the Cortical cell group showed significantly more long-rallies across time (*t* = 3.40, p = 0.0007) along with significantly fewer aces than the HEK control and Media control groups at the second timepoint (Table S2).

**Fig. S5.**
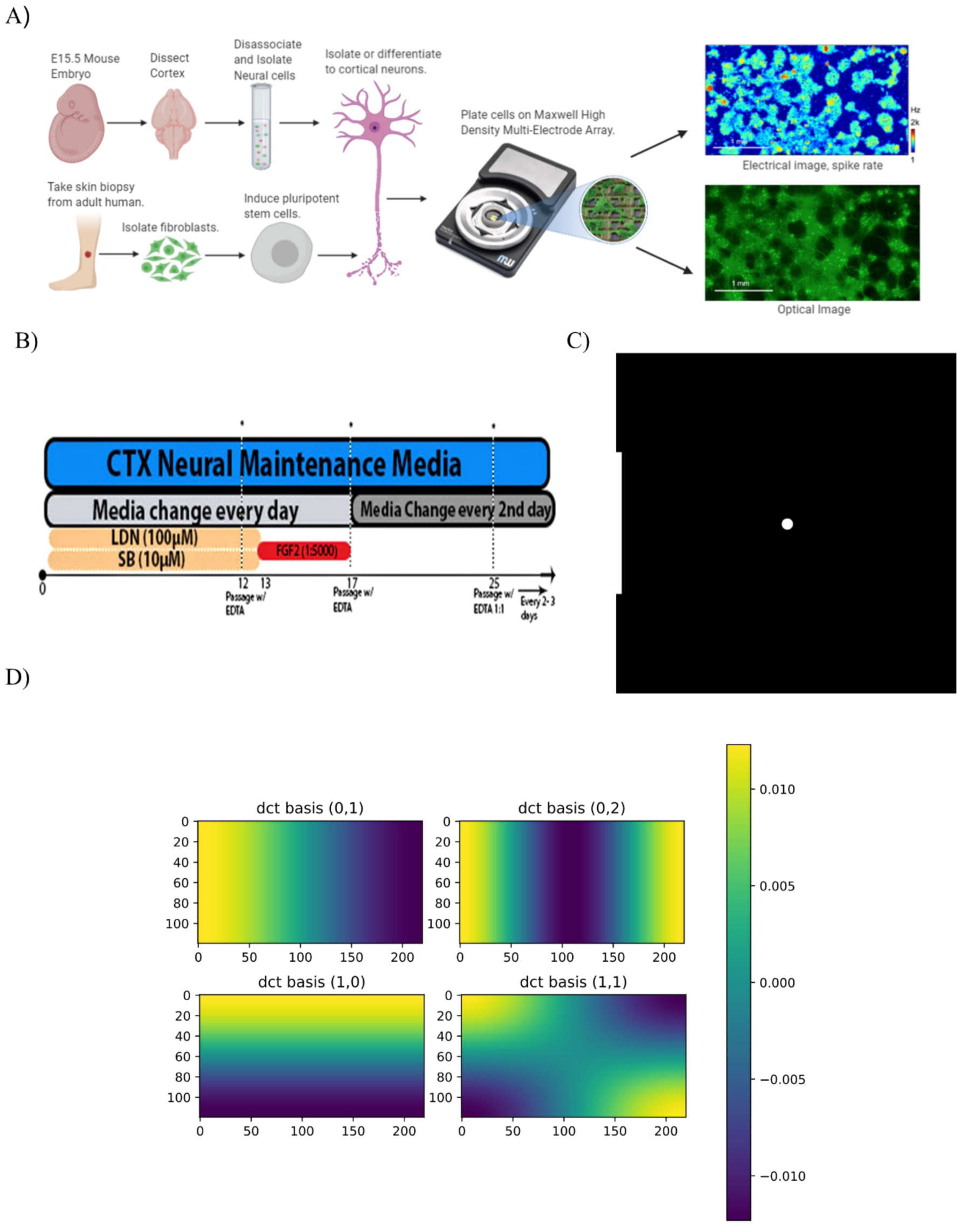
Key methods used in this study. **A)** Diagrammatic illustration of the core experimental setup which drove the research in this project. **B)** Illustration of Dual SMAD inhibition protocol for differentiating pluripotent cells into cortical cells. **C)** Starting position of paddle and ball as visualised in the *DishBrain* platform. From the perspective of the neural cultures, it is more accurate to imagine that they view this world from the perspective of the paddle looking at the ball opposed to top-down as presented here. **D)** The Discrete Cosine Transformation (DCT) Basis functions used to summarise the symmetry of spontaneous electrophysiological activity.

**Table S1.** Multivariate statistical tests and all results for tests done.

| Figure | Panel | Parameters | Source | DF1 | DF2 | MS | F | p-value | np2 | Method |
| --- | --- | --- | --- | --- | --- | --- | --- | --- | --- | --- |
| 5 | D | Average Rally Length | Group - all | 1 | 845 | 0.305 | 10.381 | 0.001 | 0.012 | ANOVA |
|  |  |  | Half - all | 2 | 845 | 0.446 | 15.172 | 0.000 | 0.035 | ANOVA |
|  |  |  | Interaction - all | 2 | 845 | 0.078 | 2.646 | 0.072 | 0.006 | ANOVA |
| 6 | B | Average Rally Length | Group - all | 4 | 394 | 0.297 | 3.330 | 0.011 | 0.033 | RM ANOVA |
|  |  |  | half - all | 1 | 394 | 9.208 | 98.908 | 0.000 | 0.201 | RM ANOVA |
|  |  |  | Interaction - all | 4 | 394 | 2.020 | 21.696 | 0.000 | 0.181 | RM ANOVA |
|  |  |  | Group – time 1 | 4 | 394 | 0.698 | 7.031 | 0.000 | 0.067 | ANOVA |
|  |  |  | Group – time 2 | 4 | 394 | 1.619 | 19.519 | 0.000 | 0.165 | ANOVA |
|  | C | % Aces | Group - all | 4 | 394 | 0.081 | 9.284 | 0.000 | 0.086 | RM ANOVA |
|  |  |  | half - all | 1 | 394 | 0.131 | 16.509 | 0.000 | 0.040 | RM ANOVA |
|  |  |  | Interaction - all | 4 | 394 | 0.058 | 7.295 | 0.000 | 0.069 | RM ANOVA |
|  |  |  | Group – time 1 | 4 | 394 | 0.044 | 4.143 | 0.003 | 0.040 | ANOVA |
|  |  |  | Group – time 2 | 4 | 394 | 0.095 | 15.583 | 0.000 | 0.137 | ANOVA |
|  | D | % Long Rally | Group - all | 4 | 394 | 0.017 | 4.767 | 0.001 | 0.046 | RM ANOVA |
|  |  |  | half - all | 1 | 394 | 0.206 | 59.746 | 0.000 | 0.132 | RM ANOVA |
|  |  |  | Interaction - all | 4 | 394 | 0.047 | 13.531 | 0.000 | 0.121 | RM ANOVA |
|  |  |  | Group – time 1 | 4 | 394 | 0.046 | 10.191 | 0.000 | 0.094 | ANOVA |
|  |  |  | Group – time 2 | 4 | 394 | 0.017 | 6.928 | 0.000 | 0.066 | ANOVA |
| 7 | C | Average Rally Length | Group - all | 2 | 353 | 20740 | 4.721 | 0.000 | 0.026 | RM ANOVA |
|  |  |  | half - all | 1 | 353 | 33440 | 16.577 | 0.000 | 0.045 | RM ANOVA |
|  |  |  | Interaction - all | 2 | 353 | 25812 | 12.795 | 0.000 | 0.068 | RM ANOVA |
|  |  |  | Group – time 1 | 2 | 483 | 7559 | 2.181 | 0.114 | 0.009 | ANOVA |
|  |  |  | Group – time 2 | 2 | 483 | 53943 | 20.507 | 0.000 | 0.078 | ANOVA |
|  | D | % Change Average Rally Length vs. Rest | Group - all | 2 | 164 | 49314 | 7.674 | 0.001 | 0.086 | RM ANOVA |
|  |  |  | Test-day - all | 2 | 328 | 16.115 | 0.037 | 0.963 | 0.000 | RM ANOVA |
|  |  |  | Interaction - all | 4 | 328 | 908.448 | 2.100 | 0.081 | 0.025 | RM ANOVA |
|  | E | % Ace vs Rest | Group - all | 2 | 353 | 19992 | 6.511 | 0.002 | 0.036 | RM ANOVA |
|  |  |  | half - all | 1 | 353 | 42.70 | 0.646 | 0.422 | 0.002 | RM ANOVA |
|  |  |  | Interaction - all | 2 | 353 | 549.025 | 8.308 | 0.000 | 0.045 | RM ANOVA |
|  |  |  | Group – time 1 | 2 | 483 | 453.464 | 2.181 | 0.127 | 0.008 | ANOVA |
|  |  |  | Group – time 2 | 2 | 483 | 2906 | 18.096 | 0.000 | 0.070 | ANOVA |
|  | F | % Ace vs Rest | Group - all | 2 | 164 | 2683 | 12.125 | 0.000 | 0.129 | RM ANOVA |
|  |  |  | Test-day - all | 2 | 328 | 110.546 | 0.971 | 0.380 | 0.006 | RM ANOVA |
|  |  |  | Interaction - all | 4 | 328 | 180.459 | 1.585 | 0.178 | 0.019 | RM ANOVA |
|  | G | % Long-Rally vs Rest | Group - all | 2 | 353 | 52.007 | 0.650 | 0.523 | 0.004 | RM ANOVA |
|  |  |  | half - all | 1 | 353 | 1089.865 | 29.932 | 0.000 | 0.078 | RM ANOVA |
|  |  |  | Interaction - all | 2 | 353 | 436.936 | 12.000 | 0.000 | 0.064 | RM ANOVA |
|  |  |  | Group – time 1 | 2 | 483 | 617.708 | 8.513 | 0.000 | 0.034 | ANOVA |
|  |  |  | Group – time 2 | 2 | 483 | 162.9<br>34 | 3.219 | 0.041 | 0.013 | ANOVA |
|  | H | % Long-Rally vs Rest | Group - all | 2 | 164 | 154.4<br>46 | 0.490 | 0.614 | 0.006 | RM ANOVA |
|  |  |  | Test-day - all | 2 | 328 | 21.67<br>8 | -0.244 | 1.000 | 0.00 | RM ANOVA |
|  |  |  | Interaction - all | 4 | 328 | 118.7<br>79 | -1.336 | 1.000 | 0.00 | RM ANOVA |
| S4 | A | % Change Average Rally Length vs. Rest | Group - all | 2 | 237 | 13897 | 3.052 | 0.049 | 0.02 | RM ANOVA |
|  |  |  | half - all | 1 | 237 | 63857 | 27.008 | 0.000 | 0.102 | RM ANOVA |
|  |  |  | Interaction - all | 2 | 237 | 13814 | 5.843 | 0.003 | 0.047 | RM ANOVA |
|  |  |  | Group – time 1 | 2 | 443 | 1389 | 0.405 | 0.667 | 0.002 | ANOVA |
|  |  |  | Group – time 2 | 2 | 442 | 42205 | 14.107 | 0.000 | 0.060 | ANOVA |
|  | B | % Ace vs Rest | Group - all | 2 | 237 | 1956 | 10.036 | 0.000 | 0.078 | RM ANOVA |
|  |  |  | half - all | 1 | 237 | 378 | 10.036 | 0.010 | 0.028 | RM ANOVA |
|  |  |  | Interaction - all | 2 | 237 | 258 | 4.596 | 0.011 | 0.037 | RM ANOVA |
|  |  |  | Group – time 1 | 2 | 443 | 844 | 5.060 | 0.007 | 0.022 | ANOVA |
|  |  |  | Group – time 2 | 2 | 442 | 2828 | 26.297 | 0.000 | 0.106 | ANOVA |
|  | C | % Long-rallies vs Rest | Group - all | 2 | 237 | 47.90 | 0.791 | 0.454 | 0.007 | ANOVA |
|  |  |  | half - all | 1 | 237 | 1507 | 33.155 | 0.000 | 0.123 | RM ANOVA |
|  |  |  | Interaction - all | 2 | 237 | 344 | 7.585 | 0.001 | 0.060 | RM ANOVA |
|  |  |  | Group – time 1 | 2 | 443 | 425 | 6.063 | 0.003 | 0.027 | RM ANOVA |
|  |  |  | Group – time 2 | 2 | 442 | 258.7 | 6.029 | 0.003 | 0.027 | ANOVA |

**Table S2.**
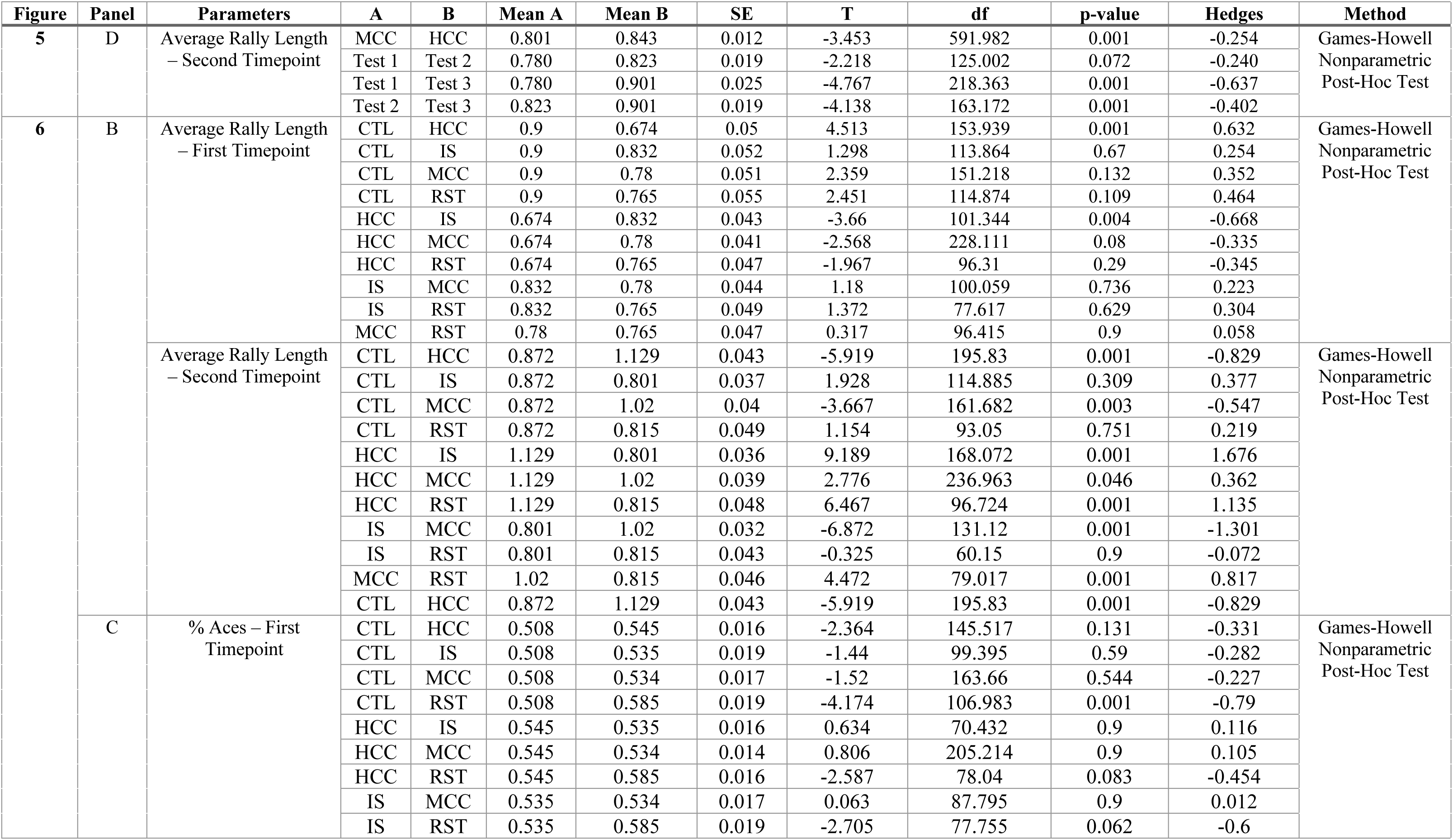
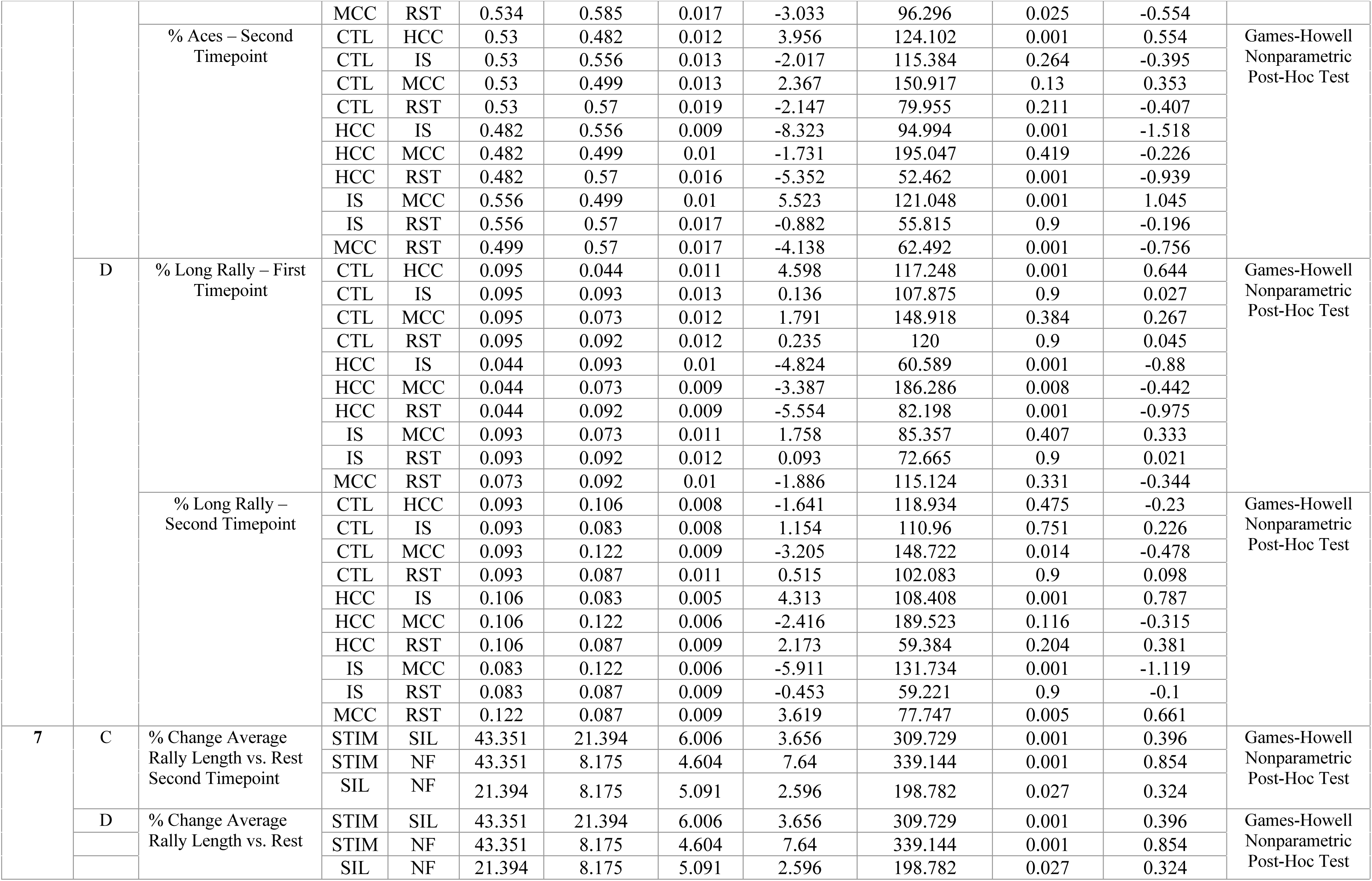
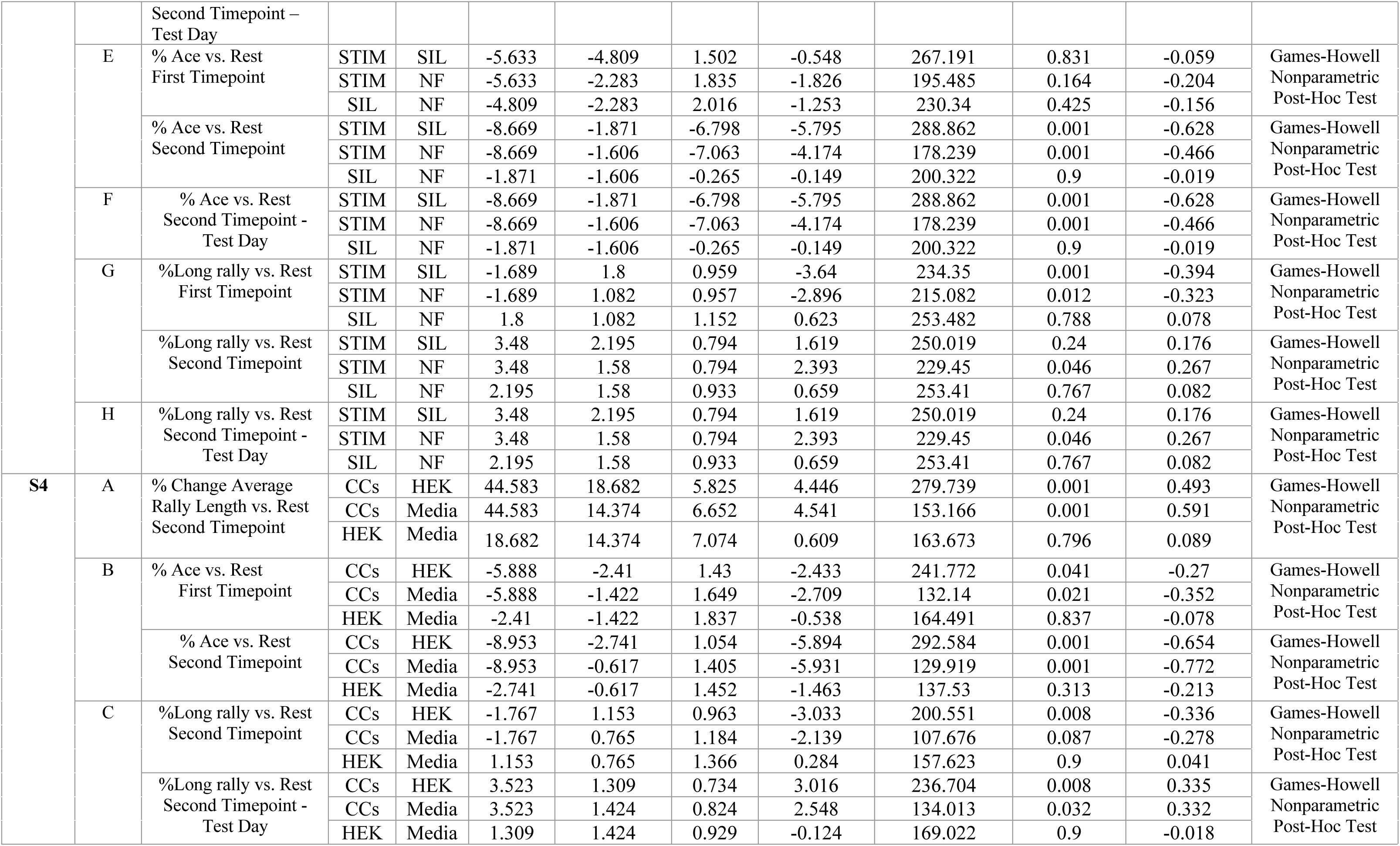
Follow up post-hoc tests for multivariate tests with exact p-values.

**Table S3.** Percentage configurations selected (in bold) by EXP3 algorithm for control and experimental groups.

| Configuration | Control<br>% | Experimental<br>% |
| --- | --- | --- |
| 0 | **22.17** | 15.51 |
| 1 | 16.16 | 16.62 |
| 2 | 13.93 | 18.50 |
| 3 | 18.49 | **19.31** |
| 4 | 12.25 | 14.69 |
| 5 | 17.01 | 15.37 |

**Movie S1.**

Representative video of a paddle being controlled by the activity of living neurons to play a simulated game of pong. It is of particular interest to note how frequently after a successful hit the paddle leads where the ball will eventually end up on the return, even before the ball hits the backwall.

**Movie S2.**

Representative video of a paddle being controlled by the activity of living neurons to play a simulated game of pong in the SpikeStream interactive visualizer. This is also available live in real time from any active culture in the *DishBrain* system.

